# Genomic hallmarks of parasexual reproduction in three hybrid groups of the human pathogen *Cryptococcus neoformans*

**DOI:** 10.64898/2026.05.28.728413

**Authors:** Rahul Anand, Qinxi Ma, Diana Tamayo, Grace Paul, Nicolas Helmstetter, Sheng Sun, Zhuyun Bian, Kyung J. Kwon-Chung, Joseph Heitman, Rhys A. Farrer

## Abstract

Hybridization is a major driver of fungal evolution, yet knowledge of the molecular mechanisms underpinning hybridization and its genomic impact remain limited. Here, we analyse 197 *Cryptococcus neoformans* genomic sequences, including 13 newly sequenced strains, identifying three genetically clustered and distinct hybrid groups (H1, H2 and H3) each with unique parental origins and ecological associations. Using phylogenomics, population structure analyses, and long-read genome assemblies, we identified hybrid genomes with chromosome-wide loss of heterozygosity (LOH), inheritance of large intact parental haplotype blocks and widespread aneuploidy within and across genomes. These patterns are also observed when progeny were generated with *spo11*Δ parents, indicating these features are a result of a meiotic-independent process such as parasex. This is further underscored by discovery of haploid and near-haploid recombinants in both *spo11*Δ mutant progeny and reanalysed wild-type hybrid progeny. We hypothesise these haploid and near-haploid recombinants are generated through ploidy reduction via independent chromosome assortment because of concerted chromosomal loss, which is a key feature of the parasexual cycle. Phenotypic assays demonstrated that several hybrid isolates have diverse growth and virulence patterns, underscoring functional consequences of genome plasticity. Together, our work suggests a non-meiotic reproductive process contributes to shape the genotypic and phenotypic diversity of *Cryptococcus neoformans*.

**Significance statement:** *Cryptococcus* is responsible for approximately 180,000 deaths annually worldwide. However, the mechanisms by which *Cryptococcus* generates genotypic diversity remains understudied. Using comparative genomics, we identified three distinct groups of hybrids, defined by their ancestral inheritance. In each group, we discovered novel genomic features including uneven chromosome numbers in most hybrids, chromosome-wide loss of heterozygosity consistent with whole chromosome inheritance from a single parent, and whole-chromosome-wide haplotype inheritance. Similar genomic patterns were observed in *spo11*Δ mutant crosses. These genomic features are consistent with a parasexual form of reproduction. Our results expand our understanding of fungal reproduction and highlight new routes for the emergence of virulence and antifungal resistance.

## Introduction

Fungal species propagate through a diverse array of strategies, encompassing both sexual and asexual modes (1). Sexual reproduction occurs via meiotic and/or mitotic recombination process during outbreeding (heterothallism) or selfing (homothallism) (1).

In heterothallic fungi, mating is initiated by the recognition of compatible partners through mating type-specific peptide pheromones and receptors, followed by cell-cell fusion. Genetic exchange typically requires compatibility at the mating-type loci (*MAT***a** and *MAT*α), which can be organised as a single genomic loci (bipolar) or across multiple loci (e.g., tripolar, tetrapolar) (2). This process culminates in the formation of spores, which facilitate dispersal and long term survival through dormancy, stress resistance, and the ability to colonise new environments (3, 4).

While meiosis is the hallmark of canonical sexual reproduction, some fungi also engage in parasexual reproduction this non-meiotic process that includes genome fusion, mitotic recombination, and ploidy reduction resulting in chromosomes originating from either parent (5–7). First described by Guido Pontecorvo in *Aspergillus nidulans* (8), parasexuality has since been reported in multiple fungal phyla including other Ascomycota species such as *Candida albicans* (6, 7), *Fusarium oxysporum* (9) and *Verticillium dahlia* (10). Evidence of parasexual cycles have been found in the Basidiomycota genus *Ustilago* (11), and in the Chytridiomycota amphibian pathogen *Batrachochytrium dendrobatidis* (12).

*C. neoformans*, *C. deneoformans*, and the *C. gattii* species complex are important human pathogens, together responsible for over 180,000 deaths annually worldwide (13). The sexual cycle of the human-pathogenic fungus *Cryptococcus neoformans* was first described in 1975 by June Kwon-Chung (14). Sexual reproduction occurs between *MAT***a** and *MAT*α cells, which fuse to form dikaryotic hyphae. Mating is influenced by environmental cues such as inositol or nutrient limitation, which has been documented in V8, bird guano, and decaying wood (4, 15, 16). At the tips of the hyphae, a basidium forms in which nuclear fusion and meiotic recombination occur, giving rise to infectious basidiospores (17). Despite having both mating types, over 98% of clinical and environmental isolates of *C. neoformans* are *MAT*α (18, 19), suggesting that *C. neoformans* primarily propagates via asexually or via unisexual (same-sex *MAT*α) reproduction (18, 20). Under appropriate nutrient conditions (MS or V8 media), *MAT*α cells can produce basidiospores independently, whereas *MAT***a** cells typically do not - potentially contributing to the global predominance of the *MAT*α genotype (21). However, a recently published paper demonstrate congenic *MAT**a*** and *MATα* were both equally well able to undergo unisexual reproduction (22).

In Europe and parts of the Americas, *Cryptococcus* hybrid isolates - particularly those between serotype A *C. neoformans* (VNI) (previously known as var. *grubii*) and serotype D *C. deneoformans* (VNIV) (previously known as var. *neoformans*) account for up to 30% of clinical cases (23), while estimates from high-burden regions such as Africa and Asia suggest hybrids account for closer to 10% of cases (24). The discovery that these AD hybrids are diploid, typically having opposite mating types revealed they are produced through mating of haploid cells of the two sister species *C. neoformans* and *C. deneoformans* (25–27). While many AD hybrid strains are no longer self-fertile, a few have been identified that retain the ability to produce monokaryotic hyphae, basidia and spores. Spore dissection revealed a low (∼5%) spore germination rate, and the spores that germinated produced progeny found to be AD diploid with no evidence of meiotic reduction to the haploid state. Thus, the genetic differences between the two parental lineages appear to preclude meiotic recombination and chromosomal reassortment. These hybrids can exhibit altered phenotypes, including enhanced virulence (28), reduced antifungal susceptibility (29), and increased resistance to ultraviolet light (30), impacting epidemiology and clinical outcomes. Natural hybrids have been recovered from both clinical and environmental sources. Laboratory-generated crosses have revealed extensive genomic plasticity following hybridization including aneuploidy, chromosomal rearrangements, and LOH (24, 31, 32). However, most studies have focused on VNI × VNIV crosses, overlooking the potential for hybridization involving other *C. neoformans* lineages such as VNII, VNBI, or VNBII (33). Therefore, the broader genomic and phenotypic consequences of hybridization remain poorly understood.

To investigate the genetic diversity and evolutionary history of *C. neoformans*, we performed phenotypic profiling and whole genome sequencing for 13 isolates that were analysed alongside 184 publicly available genome sequences. We identified three distinct *Cryptococcus neoformans* hybrid groups and resolved their parental lineage origins. We uncovered diverse genomic signatures of hybridization - including significantly larger genome sizes, aneuploidy, whole chromosome LOH and F_ST_ changes, and haplotype inheritance patterns that are hallmarks of parasexual reproduction, a mode of recombination not previously documented in *Cryptococcus*. Through generating the first ever scaffold-level phased genome assemblies of 4 hybrid isolates, we reveal the genome structure of hybrid genomes, corroborating our parasex hypothesis. Next, by generating and crossing *spo11*Δ mutants of both *C. neoformans* H99 (VNI) and *C. deneoformans* JEC20 (VNIV), we recovered similar genomic patterns, suggesting a non-meiotic Spo11-independent pathway for recombination, consistent with parasex. Hybrid isolates exhibited phenotypic variation in traits such as virulence and growth rate, highlighting functional consequences of hybridization. We suggest that parasex is a key driver of genome evolution, population structure, and phenotypic diversity in *C. neoformans*.

## Results

### Phylogenetic analysis reveals three distinct hybrid groups

A nuclear phylogeny inferred from 61,284 phylogenetically informative sites across 197 *C. neoformans* isolates resolved six major monophyletic clades corresponding to *C. neoformans* lineages (serotype A: VNIa, VNIb, VNII, VNBI, VNBII) and *C. deneoformans* (serotype D: VNIV) (**Fig. 1a; Table S1**). All hybrid isolates found are diploid or aneuploid interspecies hybrids with genomic contributions from both *C. neoformans* (serotype A) and *C. deneoformans* (serotype D). Hybrid isolates were distributed across nuclear phylogeny rather than forming a single clade. However, a subset of clinical hybrid isolates formed a distinct cluster in the nuclear phylogeny. A mitochondrial phylogeny of the 197 isolates resolved three well-supported hybrid groups (**Fig. 1b**): H1 hybrids, comprised of clinical isolates, clustered with VNII/VNBI (*C. neoformans*); H2 hybrids clustered with the VNIV lineage (*C. deneoformans*) and H3 hybrids clustered with VNI (*C. neoformans*). Constructing a tanglegram between nuclear and mitochondrial phylogenies (**Fig. 1c**), reveals extensive discordance between the two topologies.

**Figure 1.**
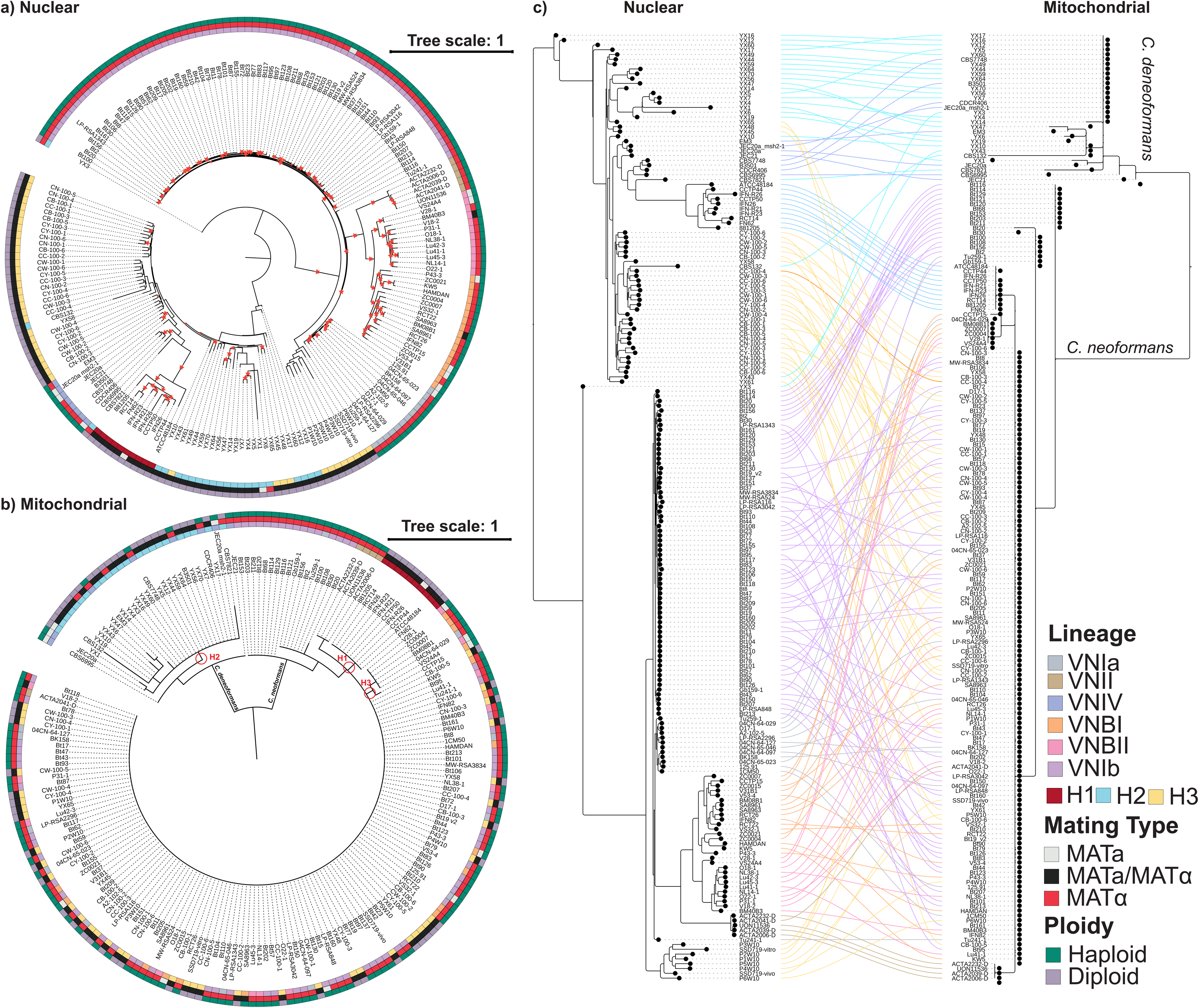
Phylogenetic analysis suggests 3 cryptococcal hybrid groups between VNBI and VNIV (H1), VNI and VNIV (H2 and H3). Phylogenetic trees for 197 *C. neoformans* isolates were constructed for the nuclear genome (**a**) and mitochondrial genome (**b**). Phylogenetically informative sites were calculated by Phylorust, where positions are covered ≥ 90% of isolates. (**c**) Tanglegram comparing the nuclear SNP phylogeny (left) and mitochondrial SNP phylogeny (right) for the same 197 isolates. The extensive mismatch in isolate ordering between the two trees indicates cytonuclear (nuclear–mitochondrial) discordance, consistent with hybridisation and subsequent recombination/backcrossing. Hybrid isolates separate into three mitochondrial-defined groups (H1–H3): H1 associates with var. grubii lineages VNII/VNBI, H3 associates with VNI (var. grubii), and H2 associates with VNIV (var. neoformans).

### Population structures of hybrids differ between the three groups

Genome-wide nucleotide diversity (π) across non-hybrid isolates was uniformly low (VNIa π = 0.002; VNIb π = 0.001; VNBI π = 0.005; VNBII π = 0.004; VNIV π = 0.005), whereas all hybrid groups display substantially elevated diversity (H1 π = 0.007; H2 π = 0.012; H3 π = 0.008) (**Fig. 2a**). Plotting π across 10 kb sliding windows reveals pronounced heterogeneity across hybrid genomes, with regions of elevated diversity with interspersed tracts of low diversity. We found that mitochondrial π is uniformly low across all hybrid isolates (**Fig. S1a)**.

**Figure 2.**
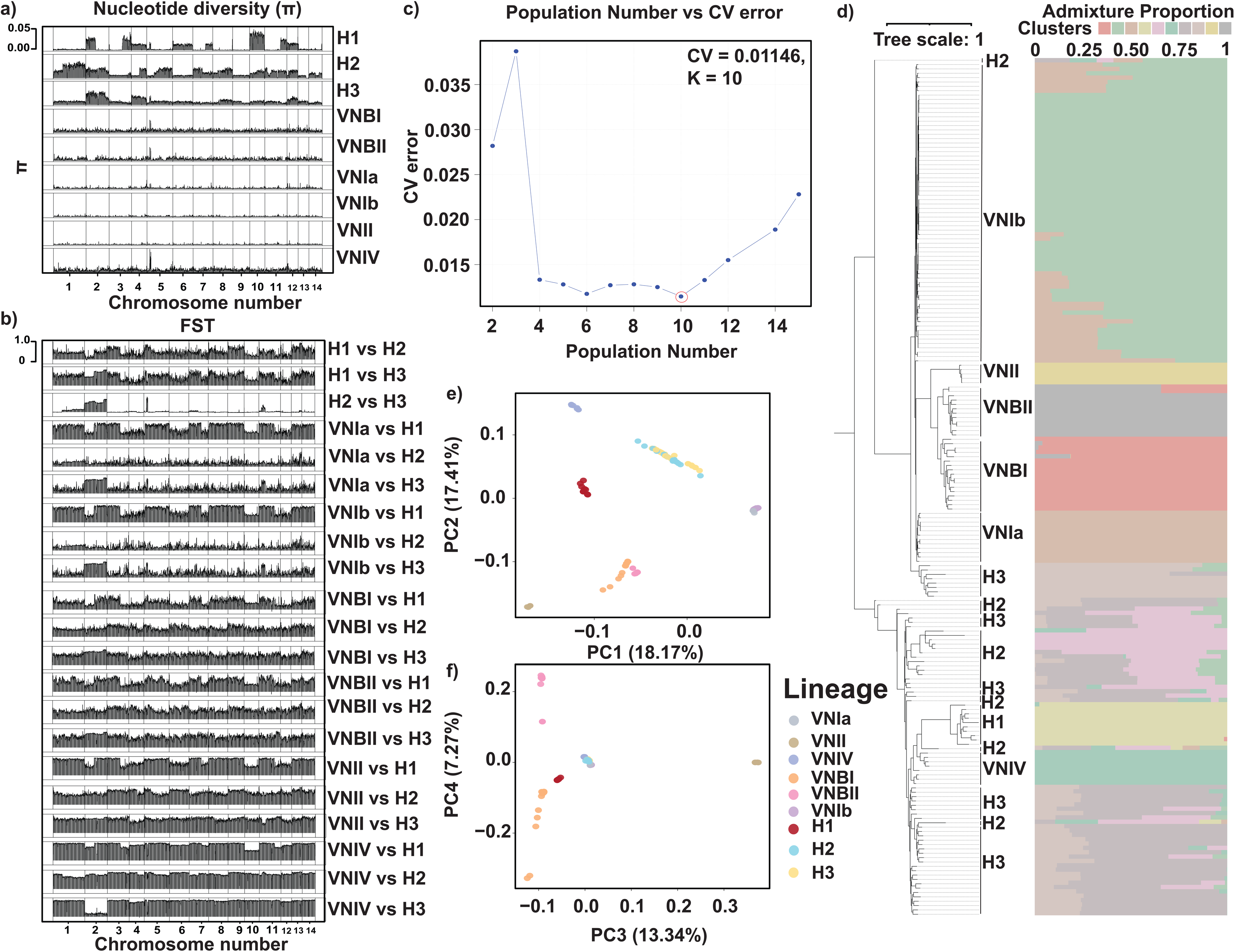
Population genomics reveals differences in nucleotide diversity, FST, and admixture between hybrid groups. **(a)** Nucleotide diversity (π) plotted across the genome for six distinct *C. neoformans* lineages and three hybrid groups. **(b)** Pairwise genetic differentiation (F_ST_) plotted across the genome, comparing the hybrid groups to each other and to the six lineages. **(c)** Cross-validation (CV) error from unsupervised ADMIXTURE analyses across K values; the lowest CV error (0.01146) occurs at K = 10. **(d)** ADMIXTURE ancestry proportions for all isolates at K = 10, showing mixed ancestry in VNIb, H2 and H3, whereas VNIa, VNII, VNBI, VNBII and H1 show little to no admixture. **(e, f)** Principal component analysis (PCA) of genomic variant sites, with isolates clustering by lineage. H1 forms a distinct cluster compared with H2 and H3. The first four principal components (PC1– PC4) together explain 56.19% of the total variance, with PC1 and PC2 shown in **(e)** and PC3 and PC4 shown in **(f)**.

Measuring nuclear divergence between the different hybrid groups, we identified minimal divergence between H2 and H3 (mean F_ST_ = 0.07), whilst both groups show high divergence from H1 (H1 vs H2 mean F_ST_ = 0.41; H1 vs H3 mean F_ST_ = 0.51) (**Fig. 2b**). In contrast, mitochondrial F_ST_ values are uniformly high when comparing hybrid groups (**Fig. S1b**) (H1 vs H2 mean F_ST_ = 0.69; H1 vs H3 mean F_ST_ = 0.96; H2 vs H3 mean F_ST_ = 0.88), including the nuclear-genetically similar H2 and H3 groups. Therefore, these hybrid groups can be distinguished by uniparental mitochondrial inheritance, with all three hybrid groups carrying diverged mitotypes, even H2 and H3, which are similar in nuclear composition.

Unsupervised ADMIXTURE analysis of nuclear SNPs identified K = 10 as the best fitting model (K = 10; CV = 0.01146) (**Fig. 2c**), resolving distinct non-hybrid lineages and revealing varying degrees of admixture among hybrid groups (**Figs. 2d, S2**). H2 and H3 show a highly admixed ancestry profiles, whereas H1 formed a distinct cluster with minimal admixture. Principal component analysis (PCA) corroborated these patterns, with H1 forming a discrete clustered whilst H2 and H3 overlapping with parental lineages (**Fig. 2e-f**). Using TriangulaR we characterised hybrid classes by plotting hybrid index against interclass heterozygosity (**Fig. S3**). Several isolates, particularly within H2 and H3, clustered near hybrid index of ≍ 0.5 and heterozygosity ∼ 1.0, consistent with being F1 hybrid progeny of recent hybridization events. However, most hybrids deviated from this expectation, demonstrating reduced heterozygosity and shifts in hybrid index towards one parental lineage.

Mitochondrial population structure was markedly simpler (K = 4; CV = 0.00395) (**Fig. S1c**). We identified a clear divide between isolates belonging to *C. deneoformans* (VNIV), which forms a single unadmixed population, and *C. neoformans* (VNI, VNBI, VNBII and VNII), which forms two populations with evidence of admixture between the two populations (**Fig. S1d**). H1 hybrids formed a distinct unadmixed mitochondrial population, while H2 hybrids fall within *C. deneoformans* and H3 hybrids fall within *C. neoformans* populations.

PCA analysis using mitochondrial SNPs corroborated this, with H1, H2 and H3 forming distinct non-overlapping positions, with H1 forming a distinct cluster, H2 clustered with VNIV (*C. deneoformans*) and H3 clustering with VNIa, VNIb and VNBI (**Fig. S1e-f**).

*MAT*α is the most common mating type (57.4%), while *MAT***a**/*MAT*α was identified in 36% of isolates, and *MAT***a** found in only 6.60% of the isolates (**Table 1**). We found that most hybrid isolates have both mating types present (*MAT***a**/*MAT*α: 93.24%; *MAT***a**: 2.70%), suggesting that unisexual hybridisation is less common (28).

**Table 1.** High prevalence of hybrids heterozygous at the *MAT* locus. Mating type percentages of each cryptococcal lineage/hybrid group.

### Hybrid genomes architectures inconsistent with a canonical meiotic origin, and shaped by aneuploidy and LOH events

Hybrid genomes show an abundance of heterozygosity across each hybrid genome, interrupted by large tracts of homozygous regions (**Figs. 3, S4**). LOH tracts ranged from localised segments to whole chromosomes, with some tracts of LOH also appearing to be localised to putative chromosome arms (e.g., on chr3 and chr11 in isolate 881205). LOH was identified in each of the three hybrid groups. However, the distribution of LOH varied substantially among isolates, with genome-wide LOH burden ranging from under 10% to over 75% within a single hybrid group (H2; **Fig. 4a**). Large contiguous LOH tracts are inconsistent with canonical meiosis alone instead being indicative of mitotic recombination such as observed as part of the parasexual cycle observed in other fungi. LOH was also not randomly distributed, with respect to the genomic context: with intergenic regions being strongly enriched, and to a lesser extent in regions flanking the genes whilst the gene body itself (5’, ORF, 3’) was depleted in LOH events occurring (**Fig. 4b**).

**Figure 3.**
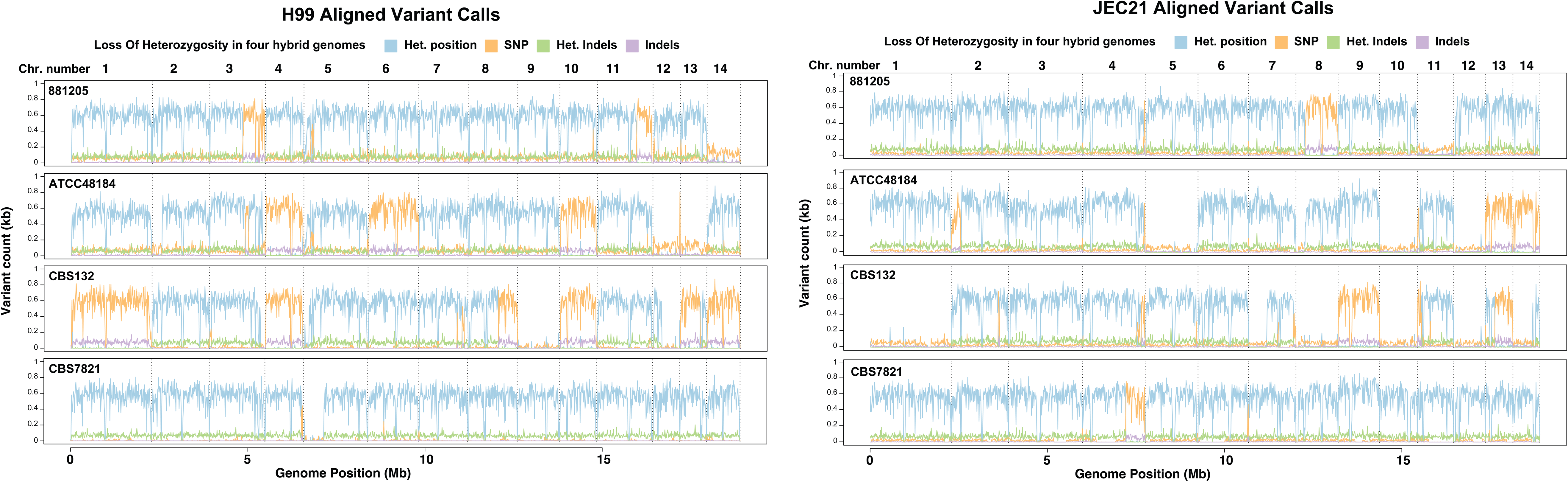
LOH and long runs of homozygosity seen throughout hybrid genome. Variant counts plotted across the genome with (blue: heterozygous positions; orange: homozygous SNPs; green: heterozygous indels; purple: homozygous indels) across 4 hybrid isolates (**a**) 881205, (**b**) ATCC48184, (**c**) CBS132 and (**d**) CBS7821 aligned to both H99 (*C. neoformans*) and JEC21 (*C. deneoformans*) reference genomes.

**Figure 4.**
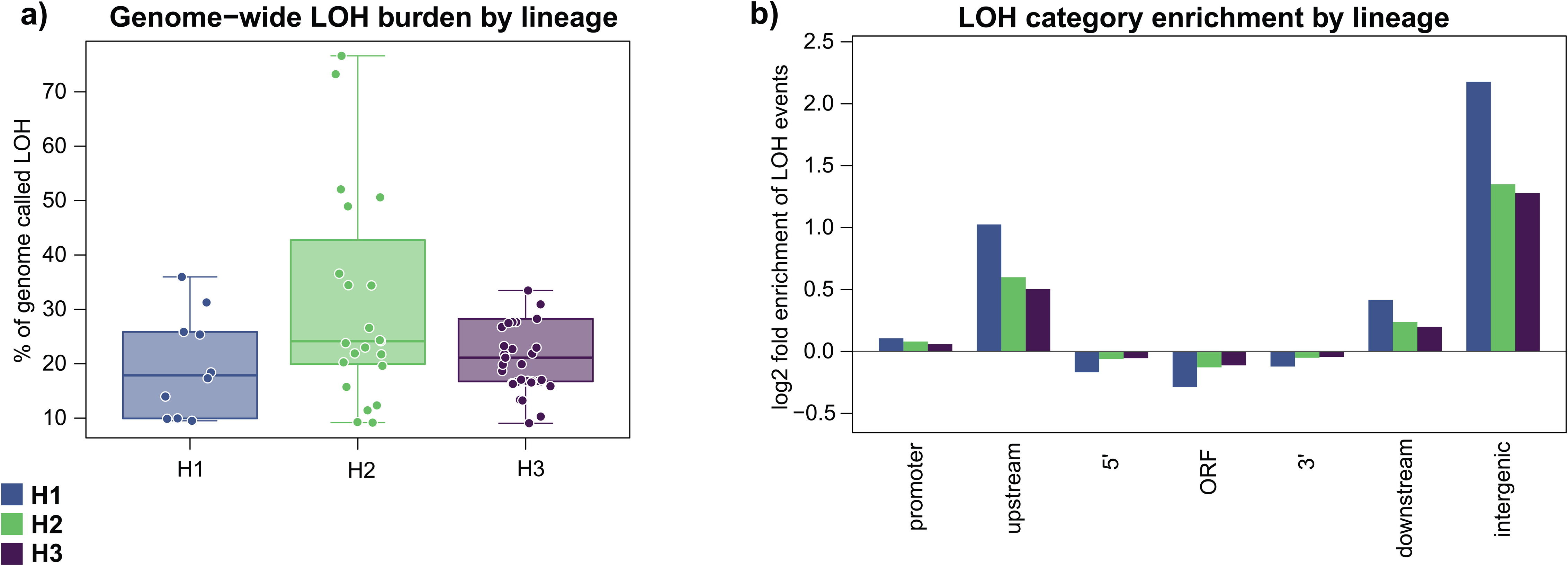
Genome-wide LOH burden and distribution of LOH relative to gene structure. LOH regions were defined as a 100 bp region where there are > 2 heterozygous positions, with (**a**) showing percentage of genome called as LOH for each isolate grouped by the 3 defined hybrid groups as box plots. (**b**) Log2fold-enrichment of LOH windows in each gene-relative category (promoter, upstream, 5’, ORF, 3’, downstream and intergenic) relative to background LOH frequency. Positive values indicate enrichment of LOH occurrences in those regions, whilst negative values indicate depletion of LOH occurrences in those regions.

To further resolve the genomic architecture of hybrid isolates, phased haplotype assemblies of four hybrid isolates were generated using long read oxford nanopore (ONT), and each separated into 2 haplotypes and a single mitochondrial sequence (**Table S2; Fig. S5**). The resulting assemblies revealed marked variation in haplotype length variation across isolates, with haplotypes ranging from 17.83 Mb (haplotype 2 CBS132) to 22.067 Mb (haplotype 2 ATCC48184).

Haplotype-specific ploidy and copy number variation is revealed through plotting depth of coverage and tally of percent of reads agreeing with the reference base across haplotypes (**Fig. 5; Fig. S6**). This divergence is further supported by low shared sequence identity between haplotypes (**Fig. S7** (34)**; Table S3**). Synteny analysis comparing two haplotypes to each other and their parental lineages (*C. neoformans* and *C. deneoformans*), reveals haplotype 1 is highly syntenic to *C. neoformans* whilst haplotype 2 is highly syntenic to *C. deneoformans* (**Fig. S8**).

**Figure 5.**
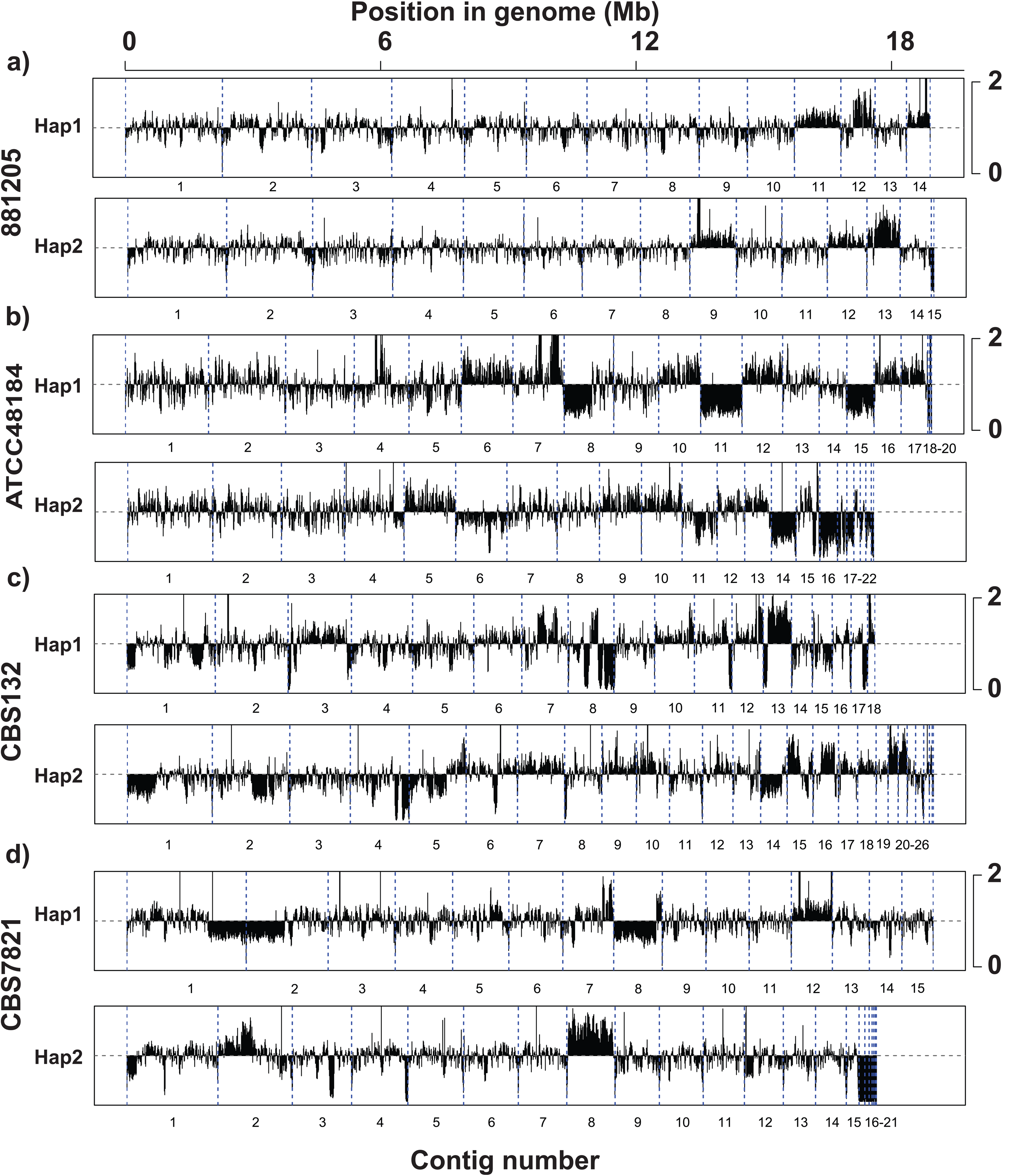
Aneuploidy and evidence of copy number variation was identified in each hybrid genome. Normalised depth of coverage plotted across haplotypes of the phased cryptococcal hybrid genome assemblies (**a**) 881205, (**b**) ATCC48184, (**c**) CBS132 and (**d**) CBS7821. Segmental aneuploidy can be seen through the genomes of all hybrid cryptococcal isolates in a “stair step” structure with regions of increased coverage flanked with normal or reduced coverage.

Phased haplotypes were assigned to their closest parental lineage by phylogenetics, across 10kb windows (**Figs. 6a, S9, Table 2**). Haplotype 2 was predominantly derived from the VNIV parent in all hybrids, whereas haplotype 1 showed variable contributions from VNIb or VNBI depending on the isolate. Parental ancestry was structured in large segmental and unbroken blocks, rather than smaller interspersed blocks. To quantify recombination, pairwise comparison of overlapping phase positions was calculated. We first used a positive control (pairwise comparison of phased *Aspergillus fumigatus*; highest percentage of crossovers identified = 0.41) (**Table S4**) and negative control (pairwise comparison of *Candida albicans*; highest percentage of crossovers identified = 0.02) (**Table S5**). The number of crossovers varied substantially between pairwise comparisons, with the highest being 393 (between 881205 and CBS7821) and the lowest being 11 (between 881205 and ATCC48184) (**Table S6**). In both cases, crossovers corresponded to a small fraction of phased positions. We classified a small number of those crossovers as potentially meiotic (crossovers that are > 1 kb apart from any other crossover and have > 2 phased positions either side of the crossover supporting the phasing). The highest number of potentially meiotic crossovers detected were between 881205 and CBS7821 (n = 21) and the lowest detected was between 881205 and ATCC48184 (n = 3). Overall, we identified potential meiotic crossover events occurring at a low frequency through hybrid genomes.

**Figure 6.**
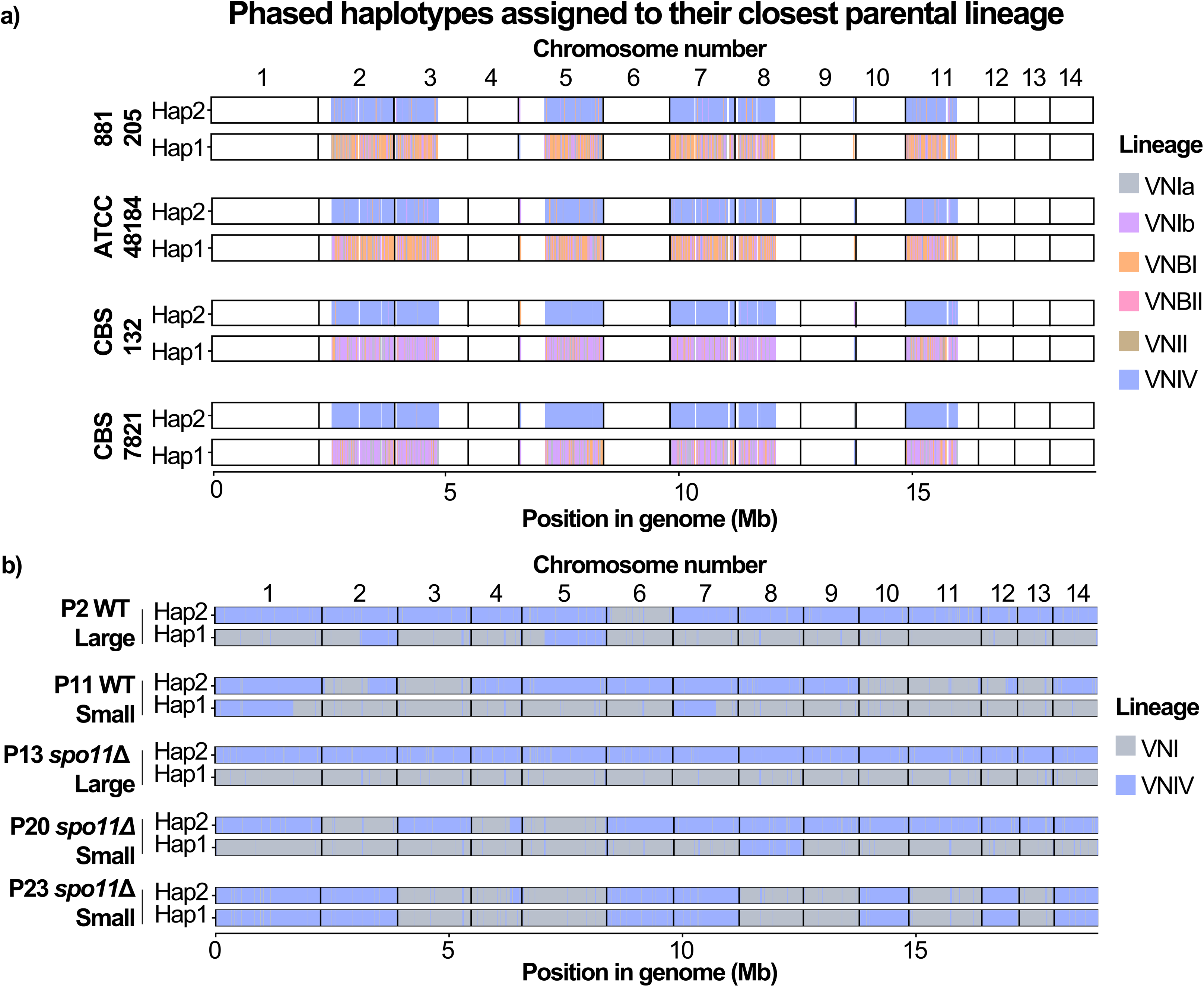
Parental-Lineage Haplotype Windows Across Hybrid Genomes. **A)** Haplotype pairs of positions that are phased in 4 hybrid isolates (881205, ATCC48184, CBS132 and CBS7821) plotted across 10kb windows, and haplotypes coloured to their closest relative (grey: VNIa, purple: VNIb, orange: VNBI, brown: VNII, blue: VNIV, white: genomic regions that are not phased in all, and therefore not directly comparable). **B)** Haplotypes pairs for lab crossed hybrids, across ‘phased in any’ sites, plotted across 10kb windows, with haplotypes coloured to their closest relative across 5 isolates (AD hybrid Progeny (P)2 Wild type (WT) Large, P11 WT Small, P13 Spo11Δ Large, P20 Spo11Δ Small and P23 Spo11Δ Small haploid progeny.

**Table 2.** Parental contributions to hybrid genomes. Haplotype clade summary of hybrid *C. neoformans* isolates, with percentage of genome inherited from each respective lineage.

### Phenotypic testing demonstrates high variability between hybrid isolates

To assess phenotypic variation, we tested the growth of eight hybrid isolates and four reference isolates (two each from *C. neoformans* (VNI) and *C. deneoformans* (VNIV) in three media types: YNBGlu, RPMI, and YPD (**Fig. S10a–c**). The *C. deneoformans* reference strain JEC21 exhibited the slowest growth in all conditions. In contrast, *C. neoformans* (H99) and *C. deneoformans* (B3501) isolates showed the fastest growth overall. Among hybrids, ATCC48184 had the highest growth in YPD and RPMI, while 881205 performed best in YNBGlu. CBS132 exhibited the slowest hybrid growth in both YPD and RPMI and was second slowest in YNBGlu. Therefore, we found no evidence for a characteristic growth rate of hybrids compared to non-hybrids. Virulence for the hybrid group isolates was evaluated using a *Galleria mellonella* infection model (**Fig. S10d**). *C. neoformans* H99 was the most virulent strain, causing 0% larval survival by day 6. In contrast, non-hybrid strains 1CM50 (*C. neoformans*), JEC21 (*C. deneoformans*), and B3501 (*C. deneoformans*) were significantly less virulent, with some larvae surviving beyond 10 days. Hybrid isolates showed wide variation in virulence: CBS132 and CBS7821 were highly virulent, resembling H99 with complete larval mortality by days 6 and 7, respectively. In contrast, 881205 and ATCC48184 exhibited attenuated virulence, with notable larval survival through the 10-day assay. Therefore, hybrid isolates do not show evidence of increased virulence, but they may have other untested advantages to non-hybrids.

### Both wild-type hybrids and *spo11*Δ lab-generated hybrids demonstrate genome restructuring consistent with parasex

To test whether the pattern of haplotype inheritance and mosaic structure of hybrid genomes is meiosis-independent, we analysed progeny derived from bilateral *spo11*Δ crosses. Spo11 is required to initiate double-stranded breaks (DSB) during meiosis, thus *spo11*Δ mutants have impaired meiosis during hybridisation. These crosses provide a means to test whether the genome patterns seen in hybrids are dependent on canonical Spo11- mediated meiosis. Interestingly, we observed a modest increase in spore germination frequency in the *spo11*Δ bilateral crosses (27%) compared to wild-type crosses (9.5%), which suggests that Spo11 is generating breaks during reproduction between *C. neoformans* and *C. deneoformans*, but these breaks are not leading to productive meiotic recombination, presumably because the DNA mismatch repair system is aborting the crossover events due to the genetic divergence in the genomes of the two parental strains. When Spo11 is mutated, these non-productive DNA breaks are no longer generated, leading to higher overall spore germination rates, although the underlying cause of this increase in germination rates cannot be precisely resolved from the present data alone.

Both wild-type and *spo11*Δ progeny showed evidence of aneuploidy and copy number variation across several chromosomes, consistent with other hybrid genomes analysed in this study (**Fig. S11**). Aneuploidy was most pronounced across small progeny, showing triploid, diploid and haploid chromosomes. Whereas large progeny showed more uniform diploidy and depth of coverage across their genomes. Strikingly, *spo11*Δ progeny 23 allele frequency distributions across the genome are broadly consistent with a largely haploid genome. Even more strikingly, half of the chromosomes are monomorphic with the *C. neoformans* H99 parental strain, whereas the other half of the chromosomes are monomorphic with the *C. deneoformans* JEC20 parent (**Fig. 6b**). Normalised depth of coverage plots also revealed some modest localised increases and decreases in coverage, indicating underlying copy number variation despite the overall haploid signal.

Plotting haplotype ancestry revealed similar proportions of parental inheritance across each of the progeny. Three isolates (P2 WT large, P13 *spo11*Δ large and P23 *spo11*Δ small) have near equal proportions of VNI and VNIV (∼50:50) (**Fig. 6b**). However, two progenies had a slight skew towards VNI (P11 WT small = 59% and P20 *spo11*Δ small = 58%). In both WT and *spo11*Δ progeny, haplotypes were inferred in large segmental blocks, as phased SNP data revealed extended genomic regions with consistent parental assignment and limited allele switching. For each of the hybrid assemblies, haplotype 1 was mostly derived from the *C. neoformans* parent while haplotype 2 was mostly derived from the *C. deneoformans* parent (**Fig. S12**). Pairwise comparison between overlapping phase positions from hybrid (and diploid) progeny revealed low numbers of crossovers (**Fig. S13**). The highest number of crossovers identified by any comparison was 13 through a comparison between P3 and P6 WT large.

### A process consistent with parasex resulting in haploid recombinant progeny

To determine if this reduction in ploidy results in haploid recombinant AD hybrids, we reanalysed sequencing data of 20 progeny reported by Kwon-Chung and Varma (2006) (35), including isolates previously classified by FACS as haploid recombinants. Plotting normalised depth of coverage (**Fig. S14a**) demonstrates copy number variations (CNVs) throughout the hybrid genomes. Plotting tally of percentage of reads agreeing with base (**Fig. S15b**) revealed 8 isolates as either predominantly or completely haploid. When plotting variants across the hybrid genomes we demonstrate LOH patterns seen in the other hybrid isolates analysed in this study (**Fig. S15**). We see LOH sometimes coinciding with a reduction in ploidy, such as that seen in chromosome 3 of isolate SB-8.

## Discussion

High-throughput sequencing has refined our understanding of *C. neoformans* phylogenomics and population structure, enabling identification of globally prevalent clades and lineage-specific variation linked to virulence (36, 37). However, *C. neoformans* hybrids have been treated as a single, poorly resolved category to date (23). Our analysis indicates this classification overlooks substantial genomic complexity. Whilst the discovery of three distinct hybrid groups (H1, H2 and H3) supports previous hypothesis stating *C. neoformans* hybrids have arisen from multiple hybridisation events (26). Here, we show that these hybrid groups differ in their parental origins, patterns of mitochondrial inheritance and population structure. A defining feature of the ancestry of each hybrid group is the uniparental mitochondrial inheritance patterns, where mitochondrial divergence was identified to be high between all three hybrid groups, including H2 and H3, which have high nuclear similarity but highly diverged mitochondria. Mitochondrial inheritance is known to contribute to fitness of fungal hybrid pathogens (38). Further investigation is now required to understand if and how differential mitochondrial inheritance impacts growth, stress response, drug resistance, and virulence.

Despite the diversity of genomic features and ancestry, inheritance and population structure *C. neoformans* hybrids have been previously grouped into a single lineage (AD or VNIII). Furthermore, studies investigating hybrids have focused primarily on VNI x VNIV crosses (33), while either neglecting or not identifying other crosses that have been described through non-whole-genome approaches, such as VNI/VNII and VNII/VNIV hybrids

(39). Findings from these studies cannot then be safely extrapolated to all *C. neoformans* hybrids, especially those of differing parental origins, such as H1 which our data associate with VNBI and VNIV parentage and show significantly divergent population structure which may impact their phenotype including their virulence. This single-lineage view underlies the evidence to date suggesting that hybrids exist in a genetically “locked in” state (28). Future investigations should consider the impact of backcrossing to parental strains and the possibility of intercrossing to generate F2 hybrids, which remains untested on fitness and virulence.

We identified *Cryptococcus* hybrids that are highly aneuploid, consistent with prior reports of genomic plasticity in *C. neoformans* hybrids (40). The specific “stair-step” copy number variation (CNV) structure we see is notable for mirroring genomic rearrangements in *Candida albicans* linked to antifungal resistance (41), raising the possibility that a comparable process occurs in *C. neoformans* hybrids. That whole-chromosome CNVs are frequent, and also supported by ploidy variation within hybrid genomes, point towards concerted chromosome loss, a mechanism known to stabilise aneuploidy and confer fitness advantages in non-meiotic parasexual fungi such as *Aspergillus nidulans* and *Candida albicans* (6, 42). We hypothesize that similar parasexual processes may drive genome stabilization and adaptation in *C. neoformans* hybrids.

Long runs of homozygosity and loss of heterozygosity (LOH) contribute to the genome plasticity seen in hybrid genomes (23, 33), in both clinical, environmental and laboratory settings. This prevalence of LOH in hybrids is difficult to reconcile with a meiotic process driving hybridisation and more consistent with parasex. Further corroborating this hypothesis is that LOH is not randomly distributed throughout hybrid genomes, and instead enriched in non-coding genomic regions whilst being depleted in coding regions. In each hybrid group there is evidence against hybrid genome restructuring occurring through a purely stochastic or group specific process, which may be driven by selection or a more targeted, perhaps epigenetic mechanism. The scale and frequency of LOH observed in all hybrids groups raises the possibility of specific environmental or host conditions that promote LOH, which has been demonstrated in other species including *C. albicans*, where *in vivo* passage is associated with LOH rates roughly 100-fold higher than *in vitro* (43). Therefore, LOH in *C. neoformans* hybrids may also contribute to within-host adaptation and increased virulence.

Varying tracts of homozygosity and heterozygosity have been observed as a key trait of parasexual reproduction in *Candida* yeast pathogens in generating diverse progeny bypassing the spore-formation process (44) and potential mechanism of rapidly acquiring antifungal drug resistance (45). We found evidence of crossovers, although haplotypes remained mostly physically segregated, paralleling patterns seen in other parasexual fungi (12). Therefore, hybrid genome structures are inconsistent with a canonical meiotic recombination pathway.

Meiosis may still be occurring in hybrid isolates of *Cryptococcus*, perhaps alongside parasex, or a parasex-like process. However, our *spo11* deficient crosses disrupt meiotic double-stranded break formation (46) making it unlikely that meiosis is responsible for hybridisation and resulting genomic restructuring observed in the resulting progeny.

Furthermore, both WT and *spo11*Δ progeny converge on the same genome structure and lack meiotic crossovers. A process producing near-identical outcomes without a functioning meiotic process points towards hybridisation within the *C. neoformans* species complex being through mitotic parasexual mechanisms, with progeny that have mostly diploid genomes that have undergone genome restructuring through LOH following hybridisation. In some rarer cases hybrids undergo concerted chromosomal loss, as seen in progeny 23, which has a haploid genome with segmental heterozygosity only at the end of chromosome 4, corresponding to the chromosomal region involved in a translocation between the H99 and JEC21 genomes (47). Progeny 23 was identified here as an F1 progeny of *spo11*Δ mutant crosses between *C. neoformans* and *C. deneoformans* (47) as a largely haploid progeny in which independent chromosomal assortment has occurred without demonstrable meiotic recombination. In previous studies, *C. neoformans* crosses with *C. deneoformans* conducted by Kwon-Chung and Varma (2006) (35) were observed to produce some progeny that were aneuploid or even near haploid cells. Our analysis of these F1 progeny similarly revealed independent chromosomal assortment without evidence of meiotic recombination. We note that these findings are in several ways analogous to what is observed during the parasexual cycle of *C. albicans* in which two diploid cells mate to produce a tetraploid that then undergoes concerted chromosome loss to return to the diploid state (44, 48) with only limited evidence of parameitoic recombination (49).

A key remaining difference between the evidence for parasex in *Cryptococcus* and in *C. albicans* is that we have not demonstrated that progeny appearing haploid regain the ability to mate, a step that would enable F2 hybrids and further backcrossing parallel to the *C. albicans* cycle. Backcrossing in H1, discussed above, is suggestive but not equivalent to this. Nor have any naturally occurring *C. neoformans* or *C. deneoformans* isolates been found carrying a full chromosome from the other species; the only reported evidence for introgression is a single ancient event (50). Further crosses from hybrid progeny that successfully generate F2 hybrids would support a parasexual cycle resembling *C. albicans*’, whereas hybrid sterility would point to something distinct closer to a sexual cycle species boundary than a functioning parasexual one.

The phenotypic variability we observe in virulence and metabolic assays raises the question of what, if anything, links it to the genomic plasticity described above. Environmental isolates CBS132 and CBS7821 stood out as the most virulent in *Galleria mellonella*, a pattern consistent with, though not proof of, environmental pressures having shaped genomic changes that enhanced pathogenic potential (51, 52); with only two isolates, we cannot rule out that this reflects isolate-specific variation unrelated to environmental origin. We hypothesize that LOH and aneuploidy, as products of parasexual reproduction, are among the mechanisms generating this variability, drawing a parallel with *Aspergillus nidulans*, where diploid isolates gain fitness under stress via mitotic recombination (53), though we have not tested whether a comparable stress-driven mechanism operates in our isolates. It remains unclear how significant these genomic differences are on virulence, but they offer insight into how hybridisation and a likely parasexual process can drive allele fixation in this human pathogen. Together, our results reveal previously unappreciated phylogenetic and population diversity arising from a non-meiotic, potentially parasexual process, diversity that may shape cryptococcal fitness and virulence.

## Materials and Methods

### DNA Sequencing

All isolates (except CBS7748 and B3501) were grown in YPD +2% glucose agar for 48h to obtain single colonies. One single colony was inoculated in YPD +2% glucose for 16h at 30°C, with shaking at 200 rpm. DNA was isolated using a combination of phenol chloroform-isoamyl-alcohol method and the Qiagen DNA mini kit (QIAmp DNA mini kit, 51304). Briefly, cells were resuspended in 200 µl of Tris EDTA buffer and 200 µl of phenol, followed by vortexing (20 sec vortex: 20 sec ice, repeated for 3 cycles). The samples were snap-frozen in liquid nitrogen, then heated to 75°C for 10 min. Next, 200 µl of chloroform:isoamylalcohol (24:1) was added and the lysate was centrifuged to separate the aqueous phase. A second chloroform:isoamylalcohol (24:1) wash was performed to remove residual phenol. The aqueous phase was precipitated with 2 volumes of absolute ethanol and incubated at -80°C for at least 2 hours. The DNA was resuspended in 400 µl of 10 mM Tris HCl pH 8.0 and treated with RNaseA (100 µg/ml) at 37°C for 30 min. DNA purification was completed using the Qiagen kit by adding 1.5 volumes of buffer AW1, followed by transfer to the DNeasy mini spin column. The column was centrifuged and washed with buffer AW2, followed by elution with 10 mM Tris HCl pH 8.0.

For isolates CBS7748 and B3501, genomic DNA was extracted from cells grown in YPD (1% yeast extract, 2% peptone, 2% glucose) and washed twice in PBS using a DNeasy Plant Pro Kit (QIAgen, #69204). Cells were lysed with 0.5 mm zirconia/silica beads in a FastPrep 24 homogenizer (MP Biomedicals) for four cycles (30s at 6m/s followed by 90s chilling on ice). All other steps followed manufacturer’s instructions.

Sequencing was performed by the Exeter Sequencing Service. Samples were prepared using the NEBNext Ultra II FS DNA Library Prep kit (NEB, #E7805) following manufacturer’s instructions. Libraries were loaded as part of a lane using XP loading on the NovaSeq 6000 using a paired-end strategy with a read length of 150 bases. All reads for these isolates have been deposited to NCBI Sequence Read Archive (SRA) under Bioproject number PRJNA1279655.

### High-molecular weight genomic DNA extraction

High-molecular weight DNA for Nanopore sequencing was extracted following Mayjonade *et al.* (2016) (54) with minor modifications. Cells were grown overnight in YPD, harvested by centrifugation, washed once with PBS, and freeze-dried overnight. The dried cell pellet was disrupted by bead-beating at 4 m/s for 10 s with 0.5 mm zirconia-silica beads using a FastPrep 24 homogenizer. A freshly prepared, pre-heated SDS-based lysis buffer (100 mM Tris-HCl pH 8.0, 50 mM EDTA pH 8.0, 0.5 M NaCl, 1% PVP40, 1% sodium metabisulfite, 5 mM DTT, 2% SDS, 200 μg/mL proteinase K and 100 μg/mL RNase A [both from 20 mg/mL stock]) was added. The sample was quickly homogenised by brief vortexing and incubated in a thermomixer at 55 °C with shaking (500 rpm, 1 h). Contaminants including proteins and polysaccharides were precipitated by adding 1/3 (vol/vol) of 5 M potassium acetate, incubating for 10 min at 4 °C, and centrifuging (8,000 × g, 10 min, 4 °C). Genomic DNA was purified from the cleared lysate using a NucleoMag® DNA Bacteria kit (Macherey-Nagel, #744310.4), according to the manufacturer’s instructions starting at step 4 (DNA binding). DNA integrity was confirmed on a 0.5% agarose gel, and purity and concentration were assessed by NanoDrop spectrophotometry and Qubit fluorometry.

Genomic DNA was sequenced by the Exeter Sequencing Service on the ONT PromethION using R10.4.1 flow cells with Kit 14 (E8.2) chemistry. Reads were basecalled and demultiplexed with Dorado using the super-accurate model <u>dna_r10.4.1_e8.2_400bps_sup@v4.3.0</u>. All reads for these isolates have been deposited to NCBI Sequence Read Archive (SRA) under Bioproject number PRJNA1322120.

### Genome assembly

ONT reads were filtered with Filtlong v0.3.1, favouring longer, higher-quality reads and downsampling to ∼100× where possible (filtlong --min_length 1000 --min_mean_q 90 --target_bases 2000000000 --length_weight 10). To enable haplotype-resolved hybrid genome assembly, parental *k*-mer profiles were generated directly from the reference assemblies of *C. neoformans* H99 and *C. deneoformans* JEC21 using **yak** (v0.1) (https://github.com/lh3/yak) with “yak count -k 31 -b 32 -o out.yak in.fasta”. Hybrid assemblies were then generated with **hifiasm** (v0.25.0) (55) using Oxford Nanopore long reads and the parental *k*-mer databases to guide competitive read binning and assembly of phase-separated haplotigs: “hifiasm --ont reads.fastq.gz -1 H99.yak -2 JEC21.yak -c 1 -d 1 -o output”. For isolates CBS132 and ATCC48184, the --trio-dual option was additionally applied to improve parental haplotype balance.

Each haplotype assembly was polished using Illumina paired-end reads and **Hapo-G** (v1.3.8) (56). To assign short reads to their respective haplotypes, Illumina reads were first mapped competitively against both haploid assemblies using **BWA-MEM2** (**57**) (v2.2.1). The combined reference was indexed, and alignments were sorted and indexed with **samtools** (v1.22). Polishing was performed independently for each haplotype using Hapo-G e.g., “hapog --genome hap1.fasta --pe1 hap1.R1.fq.gz --pe2 hap1.R2.fq.gz -o hap1_hapog -u”.

This procedure corrected residual base-level errors while preserving haplotype phasing, resulting in high-quality, polished assemblies for both parental subgenomes.

The completeness of the assemblies was assessed with **BUSCO** v6.0.0 (58) using the basidiomycota odb12 lineage dataset to search for near-universal single-copy orthologs. Phasing accuracy was assessed with **yak trioeval** using parental k-mer databases (H99, JEC21), reporting genome-wide SwitchErr% (phase switches between informative sites) and HammingErr% (sites needing relabelling for consistent phase) from phase-informative k-mers across hap1/hap2.

These Whole Genome Shotgun projects have been deposited at NCBI GenBank under the accessions JBYKLF000000000 and JBYKLG000000000 (isolate 881205), JBYKLH000000000 and JBYKLI000000000 (isolate CBS7821), JBYKLJ000000000 and JBYKLK000000000 (isolate CBS132), and BYKLL000000000 and JBYKLM000000000 (isolate ATCC 48184)

### Read alignment and variant calling

In addition to the 13 isolates from our collection, we also analysed an additional 184 isolates as obtained from the NCBI SRA, selected based on the following criteria: whole-genome Illumina paired-end sequencing and taxonomic classification as *C. neoformans*, *C. deneoformans*, or *C. neoformans × C. deneoformans* hybrids. Isolates were chosen to represent all major lineages (VNI, VNII, VNB, and VNIV). We also included the MATa isolate JEC20a, as no other representative of MATa was present in our dataset. Public isolates were sourced from 10 BioProjects: PRJEB14735, PRJEB27222, PRJNA1076182, PRJNA174540, PRJNA191162, PRJNA382844 (59), PRJNA626676, PRJNA700784, PRJNA764746 (37) and PRJNA999197 (35). PRJNA700784 was included for its focus on AD hybrid isolates. Reads obtained from the NCBI SRA databases were converted to the FASTQ format using SRA Toolkit v3.0.8.

Illumina reads were aligned to 4 reference sequences: 1) *C. neoformans* H99 CNA3 (GCA_000149245.3), 2) *C. neoformans* H99 mitochondrion (CP003834.1), 3) *C. neoformans* H99 MATα locus (AF542529.2) and 4) *C. neoformans* JEC20 MATa (AF542530.2) using the Burrows Wheeler Aligner (BWA) v0.7.17-r1188 mem(57). Alignments were converted to sorted BAMs using SAMtools v1.21(60).

Variant calling was performed as previously described (61). Briefly, duplicates were marked in the alignments, and the resulting file was sorted by coordinate order. GATK v4.1.2.0 (62) HaplotypeCaller was run in GVCF mode on each isolate individually, using the haploid or diploid ploidy flag as appropriate, to produce one GVCF per isolate. Per-isolate GVCFs were combined using GenomicsDBImport and jointly genotyped with GenotypeGVCFs to produce a single multi-sample VCF, which was then hard filtered (QD < 2.0, FS > 60.0, MQ < 40.0).

### LOH Determination, Burden and Structural Enrichment Analysis

Heterozygous variant positions were extracted from per-isolate VCF files and clustered into heterozygous regions, defined as ≥2 heterozygous positions within 100bp of one another. Each contig was tiled into 1kb windows, and the proportion of homozygous sequence in each window was calculated as one minus the fraction of the window overlapping a heterozygous region. Windows with a homozygous fraction ≥ 0.9 were called as LOH. LOH burden was calculated as the summed length of LOH windows as a percentage of the whole genome length, and isolate LOH burden was grouped by hybrid group (H1, H2, H3) for comparison.

To test whether LOH was non-randomly distributed with respect to gene structure, genes annotated in the reference GFF3 were used to classify flanking and internal regions as promoter, upstream, 5′, ORF, 3′, and downstream (each bounded by a defined size cap or the neighbouring gene, and assigned relative to each gene’s strand); any 1kb window not overlapping these regions was classified as intergenic. An expected background distribution was generated by classifying every 1kb window across the whole genome into these categories independent of isolate variant calls, giving each category’s proportion of the genome under a null of no association with LOH. For each hybrid group and category, the observed proportion of that group’s LOH windows falling into the category was compared against this expected genome-wide proportion using a 2×2 chi-squared test, and fold enrichment was calculated as the ratio of observed to expected proportion, presented as log2(fold enrichment), with enrichment positive and depletion negative relative to zero. As adjacent windows within a single LOH tract are not fully independent, fold enrichment magnitude and its consistency across hybrid groups were treated as the primary evidence of non-random distribution, rather than p-values alone.

### Phylogenetic and population genomics analysis

The sites passing filtering after variant calling of the VCF files of 197 *C. neoformans* isolates were converted to a multi-sample FASTA using Phylorust (https://github.com/rhysf/Phylorust), using default parameters (-s 1 [homozygous SNPs only], -m 4 [min depth 4], -p [percent threshold to extract sites 90]). The resulting multi-sample FASTA file with 55441 nuclear sites per isolate. The same parameters were used for VCF files that were constructed using alignments made with CP003834.1 H99 mitochondrial reference resulting in a multi-sample FASTA file with 12 mitochondrial sites per isolates. Phylogenetic trees of both nuclear and mitochondrial FASTA files were generated using RAxML v8.2.13(63) with 1000 bootstraps and generalized time-reversible (GTR) model and category (CAT) rate approximation model with GTR-Gamma distributed rates for the final evaluation of trees. A SplitsTree(64) Bio-Neighbour Joining Network analysis was made using the same nuclear sites.

The VCF files of the 197 *C. neoformans* isolates were indexed and merged using VCFtools (65), retaining all sites (including invariant) to produce an all-sites VCF suitable for population genomic summary statistics. A variant-only subset of this merged VCF was converted into ped and map format with plink 1.9 (66). Linkage disequilibrium pruning was done on the SNPs, which then was run through an unsupervised ADMIXTURE (67) analysis with random seed (settings: -haploid=”*” and -s time) for K values 1 to 15, with K = 10 having the lowest CV error. For nucleotide diversity (π) and Hudson’s F_ST_ (68) estimation via Pixy (69), using the all-sites merged VCF.

Using SAMtools v1.21 (60), BAM files were converted to SAM and mpileup format, which were used to calculate average depth of coverage and to plot normalized depth of coverage across 10 kb windows for the *C. neoformans* genomes. Normalized depth of coverage was calculated using de novo Perl scripts, where depth at each position was divided by the isolate’s average depth of coverage. Likewise, numbers of heterozygous and homozygous SNPs were counted across 10 kb windows and plotted across the genome.

VCF and BAM files were inputted into the HaplotypeTools (12) pipeline to phase VCF files and find parental inheritance of haplotypes of *C. neoformans* genomes. Briefly the script HaplotypeTools.pl was used to phase genomes, with resulting phased VCF files inputted into the utility script VCF_phased_to_PIA.pl with setting -p 2 to find most informative phased sites in that sample and FASTA sequences of haplotype blocks/pairs using the HaplotypeTools utility script VCF_phased_and_PIA_to_FASTA.pl. HaplotypePlacer.pl was used to find the nearest relative of each hybrids by iteratively constructing phylogenetic trees using FastTree(70) to find nearest relative for each haplotype.

### Antifungal resistance assay and growth curves

All 13 strains were grown overnight in YPD at 30 °C with shaking. Cells were collected by centrifugation at 4,000 rpm for 5 min, washed twice with PBS, and adjusted to OD_600_=0.02 in YPD, RPMI1640 (supplemented with 2% glucose and 0.165 M 3-N-morpholinopropanesulfonic acid (MOPS), pH=7), or YNBGlu (2% glucose, 0.65% yeast nitrogen base without amino acids). OD_600_ was read hourly by a Tecan M-Plex plate reader at 30 °C for 48 h. The mean of four independent measurements was determined for each strain. Area under the curve (AUC) was calculated by using GraphPad Prism (version 10.2.1).

Virulence assay was performed in the invertebrate model *Galleria mellonella*. *C. neoformans* strains were grown overnight in YPD at 30 °C with shaking. Cells were inoculated to YPD from overnight cultures were inoculated to 10 ml YPD to OD_600_ = 0.2 and grown for 4 h at 30 °C. Cells were collected by centrifugation at 4,000 rpm for 5 min, washed twice with PBS, and adjusted to 1×10^7^ cells/ml in PBS. Each larva was injected with 10 µl of fungal cell suspension (10^5^ fungal cells) containing 10 µg ampicillin and 3 µg kanamycin through the last pro-leg on the left. PBS containing antibiotics was included as control. 20 larvae were infected per strain. The larvae were then incubated at 37°C in the dark and survival quantified for 10 days post-infection.

### Crosses involving *spo11*Δ deletion mutants and analyses of random spore

To analyse the effect of Spo11 during hybridization between *C. neoformans* and *C. deneoformans*, we conducted two types of crosses: H99 (*C. neoformans*, wild type) × JEC20 (*C. deneoformans*, wild type) and H99 *spo11*Δ::*NAT* × JEC20 *spo11*Δ::*NAT*, and recovered random spores by microdissection following protocols published previously (71, 72). Spore germination rates were calculated and then 6 large and 6 small colonies from each cross were patched, from which genomic DNA was purified and sequenced at the Duke Sequencing and Genomic Technologies core facility (https://genome.duke.edu), on a Novaseq X platform with 250 bp paired-end option. All reads for these isolates have been deposited to NCBI Sequence Read Archive (SRA) under Bioproject number PRJNA1471625.

## Acknowledgements

RAF was supported by a Wellcome Career Development Award (225303/Z/22/Z). Efforts by S.S., Z.B., and J.H. were supported by NIH/NIAID R01 grants AI039115-28 and AI050113-20 (to J.H.). J.H. is also co-director and fellow of CIFAR program Fungal Kingdom: Threats & Opportunities. This project utilised equipment funded by the UK Medical Research Council (MRC) Clinical Research Infrastructure Initiative (award number MR/M008924/1)

