## Supplemental information for "Genomic hallmarks of parasexual reproduction in three hybrid groups of the human pathogen *Cryptococcus neoformans*"

#### Supplementary Figures

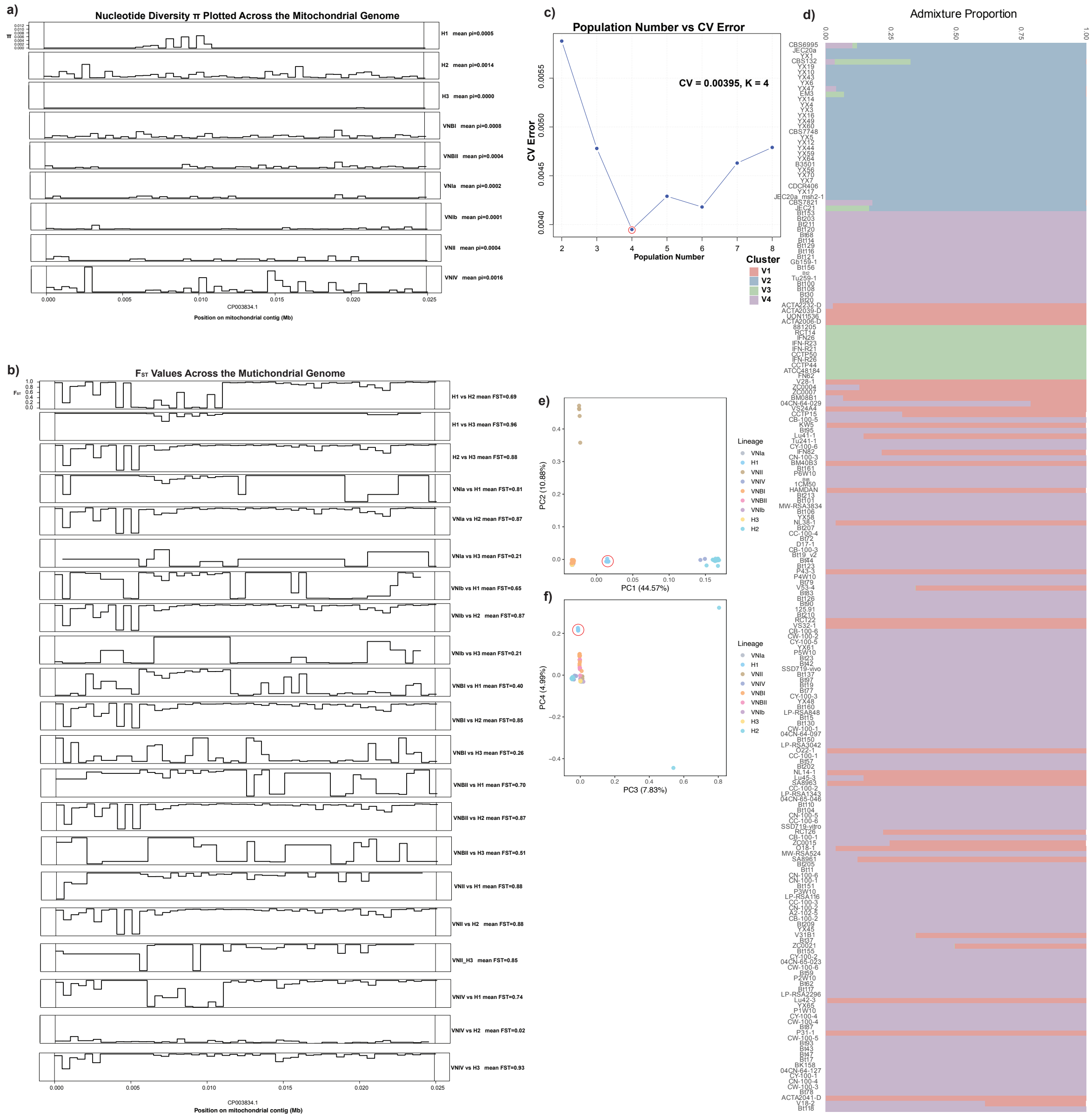


**Figure S1 Mitochondrial population genetic analysis. (a)** Nucleotide diversity ($\pi$) plotted across the mitochondrial genome. **(b)** Hudson $F_{ST}$ values plotted across the mitochondrial genome. **(c)** Unsupervised admixture analysis using mitochondrial SNP sites revealed a population size of 4 (K=11; CV error = 0.00395). **(d)** Admixture plot, show 4 distinct largely unadmixed mitochondrial populations, with var. *grubii* consisting of two populations, whilst var. *neoformans* consists of 1 population. Interestingly H1 hybrids show a distinct mitochondrial population, with no evidence of admixture. **(e,f)**, Principal component analysis (PCA) of mitochondrial SNPs similarly demonstrate H1 hybrids forming a distinct cluster, apart from the rest of the hybrids and non-hybrid lineages.

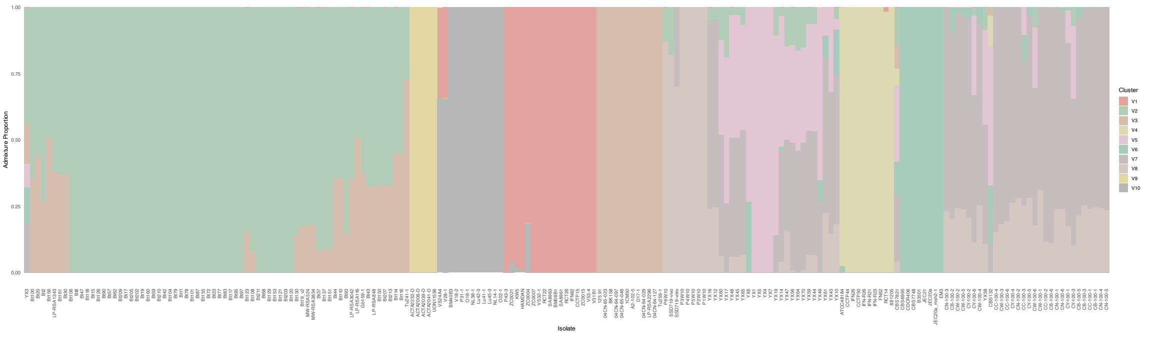


**Figure S2** ADMIXTURE plot with labels from the nuclear genomes of 197 isolates of C. neoformans, K=10.

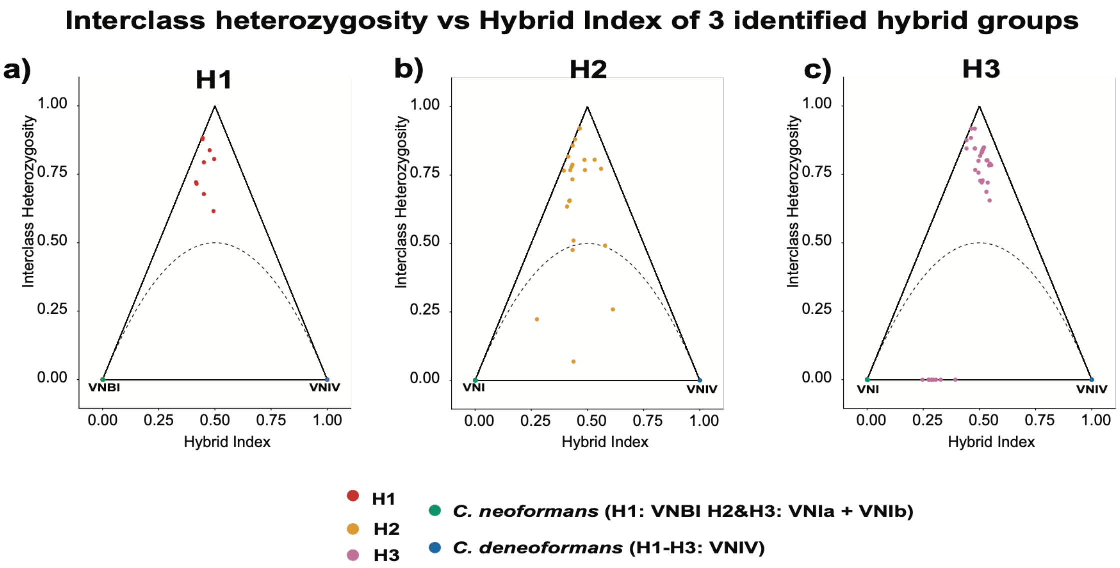


**Figure S3 TrianglulaR plots of hybrid index versus interclass heterozygosity indicate predominantly F1 hybrids.** Plots generated using triangulaR based on ancestry-informative markers (AIMs) (1) show hybrid isolates relative to parental lineages. (**a**) H1 (VNBI x VNIV); (**b**) H2 (VNI x VNIV) and (**c**) H3 (VNI x VNIV). Isolates clustering near hybrid index ≈ 0.5 with a high level of heterozygosity ~ 1.0 are consistent with F1 hybrids, whereas a lower heterozygosity suggests isolates belonging to later generation hybrids.

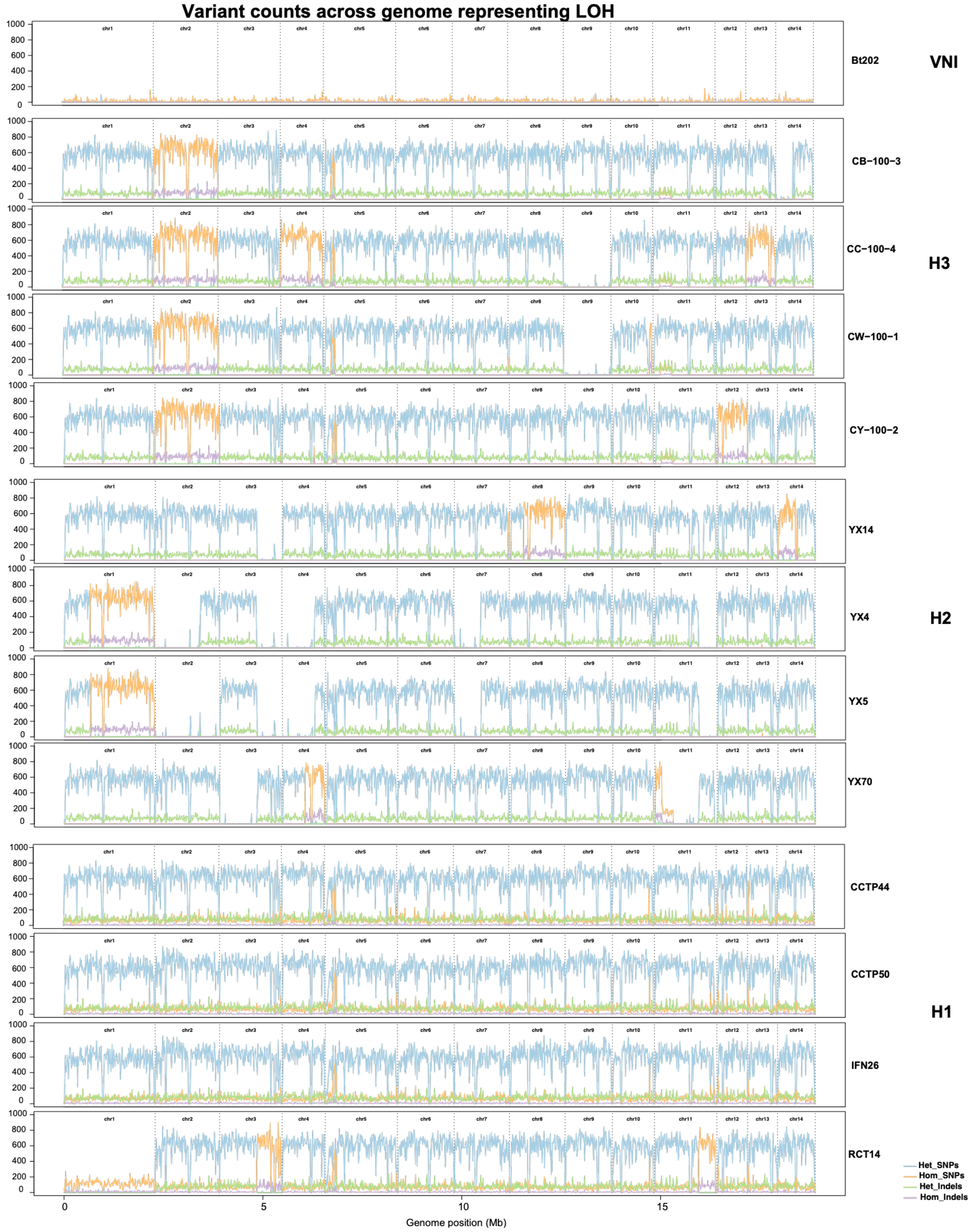


**Figure S4 LOH observed across hybrid genomes.** Variants plotted across genome with (blue: heterozygous positions; orange: homozygous SNPs; green: heterozygous indels; purple: homozygous indels) across *C. neoformans* genomes, including 12 hybrids from each representative hybrid group identified in our analysis (1: VNI; 4: H1; 4: H2; 4: H3).

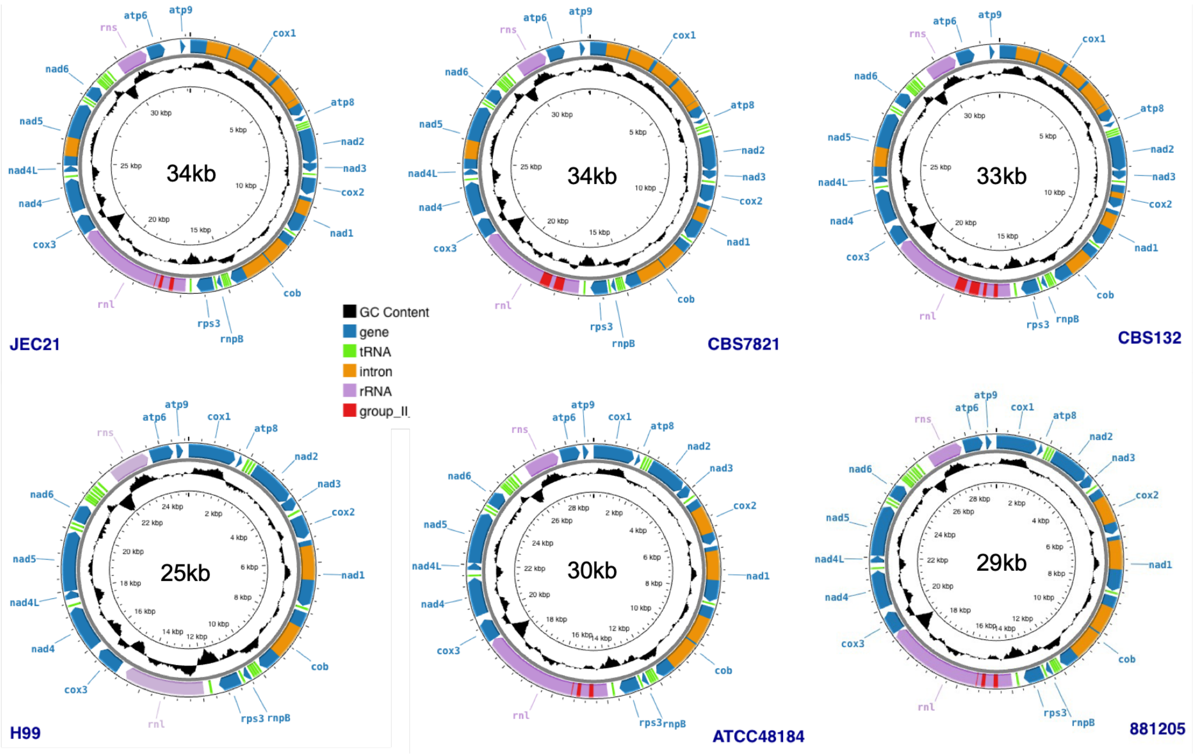


**Figure S5** Mitochondrial maps.

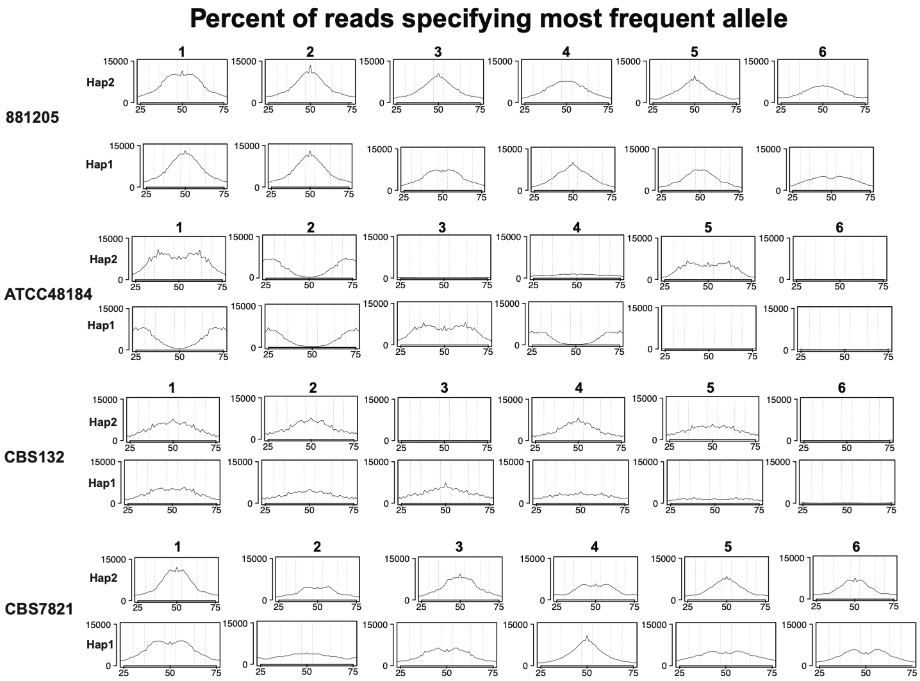


**Figure S6 Ploidy variation seen across phased long read hybrid assemblies.** Alleles frequencies (percent of reads to agreeing with the reference base) plotted from 25% agree to 75% disagree. Red-dotted lines that indicate the highest support for diploidy are at 46-53% whilst lines at 30-36% and 63-69% show greatest support for triploidy. These frequencies plotted from contigs 1-6 with 4 *Cryptococcal* hybrid genomes.

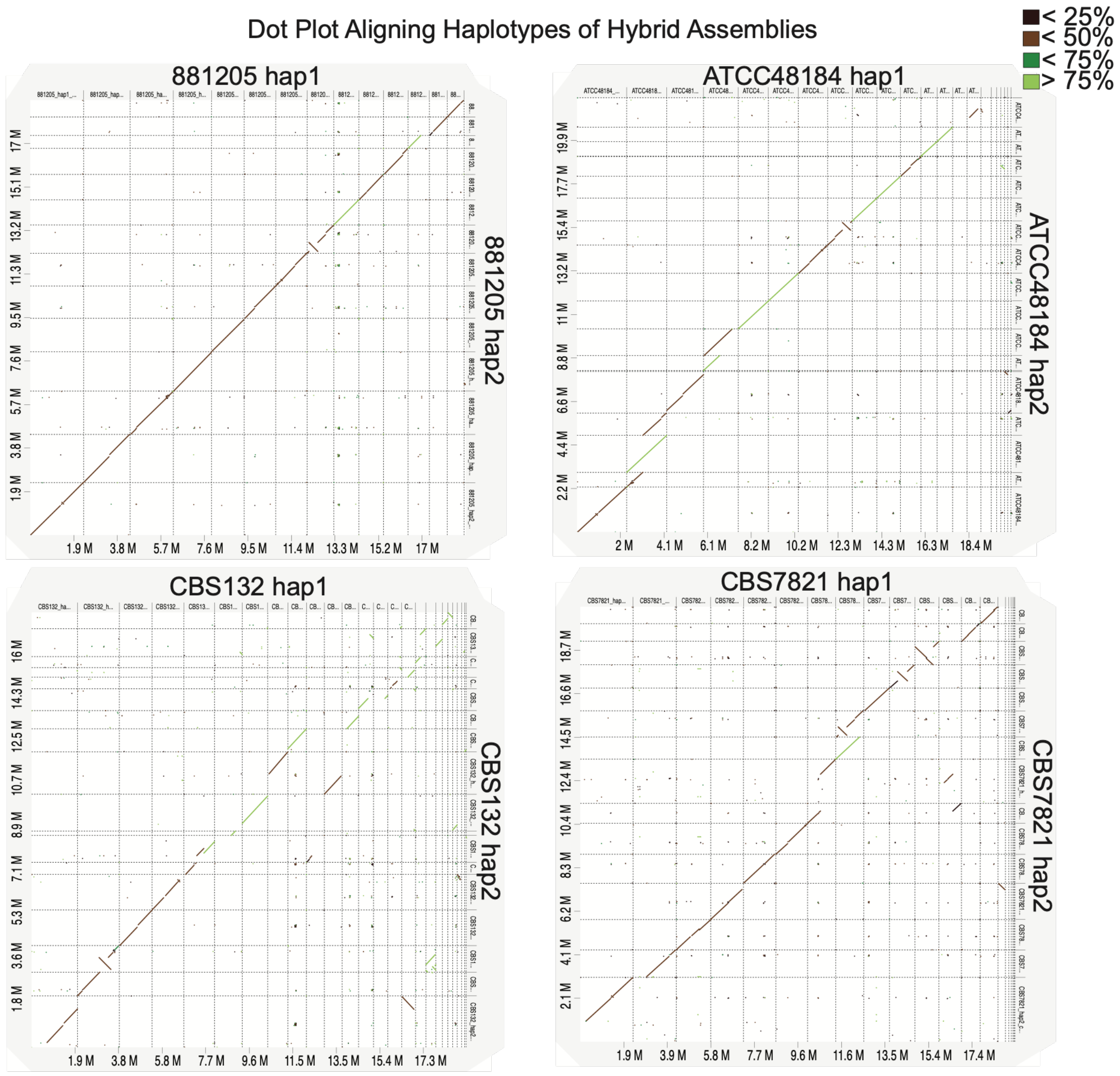


**Figure S7 Dot plots reveal divergent nature of hybrid haplotype sequences.** Dot plot aligning haplotype 1 and 2 using minimap2 from phased ONT long-read hybrid assemblies and dot plots generated through D-Genies (Cabanettes, Floréal, and Christophe Klopp. "D-GENIES: dot plot large genomes in an interactive, efficient and simple way." *PeerJ* vol. 6 e4958. 4 Jun. 2018, doi:10.7717/peerj.4958). Black is sequences with < 25% identity; brown < 50%; dark green < 75% identity; light green > 75% identity.

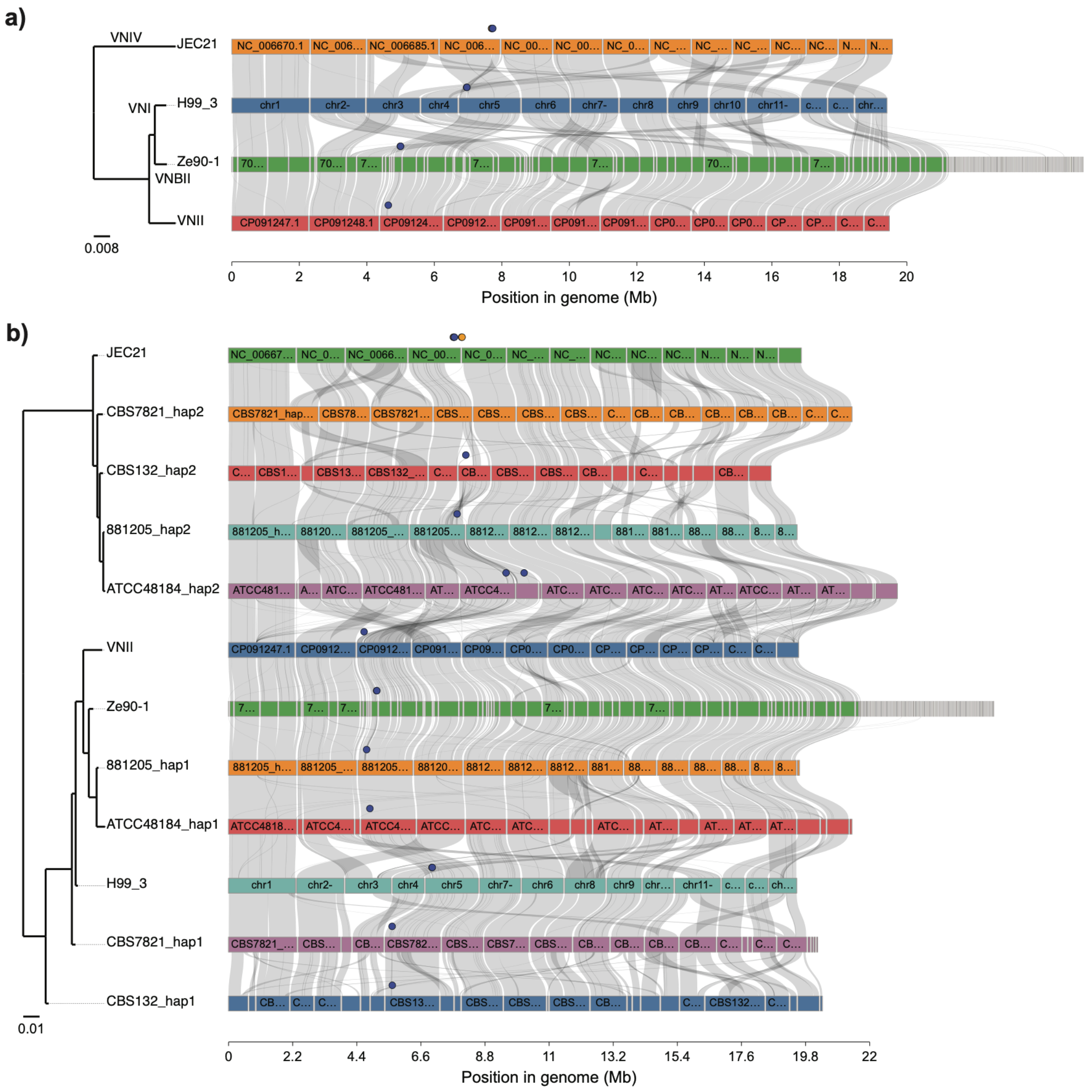


**Figure S8. Synteny between the 2 haplotypes of hybrid assemblies and reference genome assemblies, showing the location of the MAT locus (blue circle) and STE when not flanking the MAT locus (orange circle). A)** Synteny between reference genome assemblies. **B)** Synteny between haplotypes of phased assemblies and parental reference genomes of H99_3 (*C. neoformans*) and JEC21 (*C. deneoformans*).

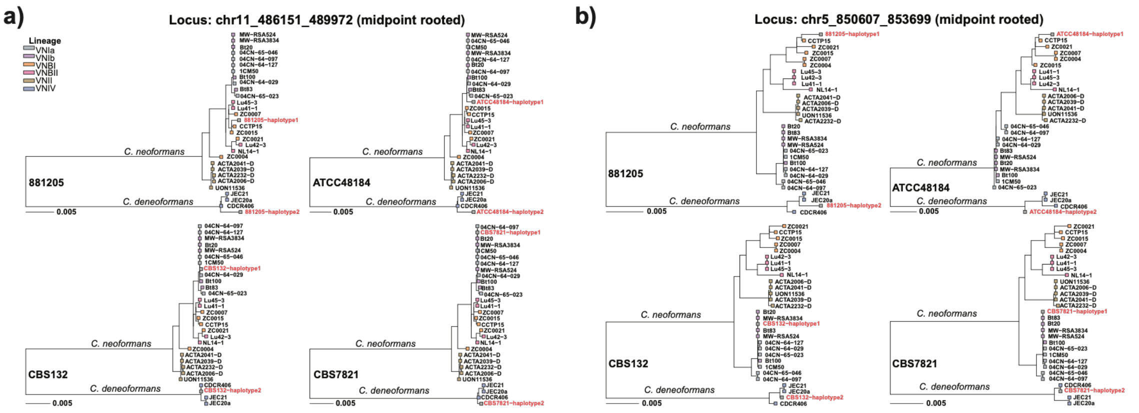


**Figure S9 Phylogenetic trees of largest haplotypes reveal segregation of parental haplotypes.** Phylogenetic tree of 2 of the largest haplotypes found (**a**) chromosome 11, locus 486151-489972 which is 3821 bp long and (**b**) chromosome 5, locus 850607-853699 which is 3092 bp long.

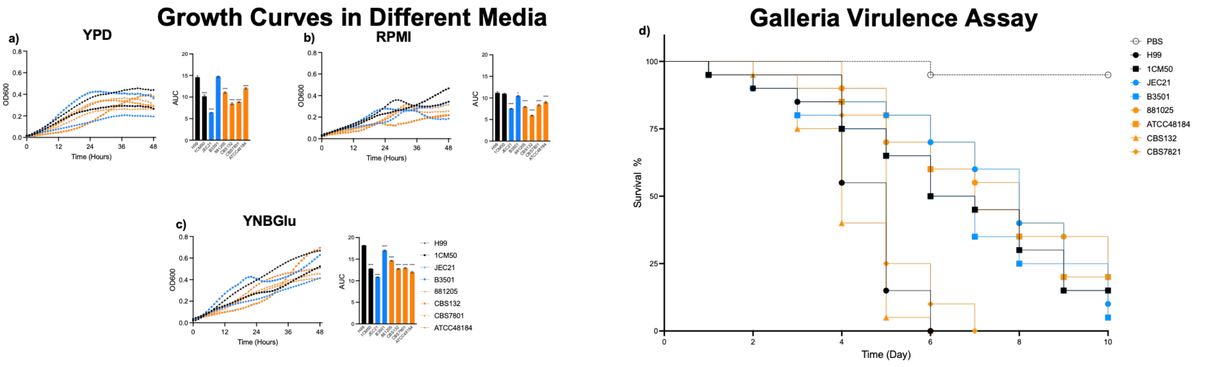


**Figure S10 Phenotypic and virulence profiles for hybrid isolates.** Phenotypic assays assess growth and virulence in eight *C. neoformans* isolates, including four hybrids. Black = VNI, blue = VNIV, orange = hybrids. (**a-c**) Growth curves and area under the curve (AUC) comparisons are shown for YPD, RPMI, and YNB+glucose conditions. * indicates difference compared to *C. neoformans* H99 (**d**) Virulence was evaluated in *Galleria mellonella*, with survival plotted across time.

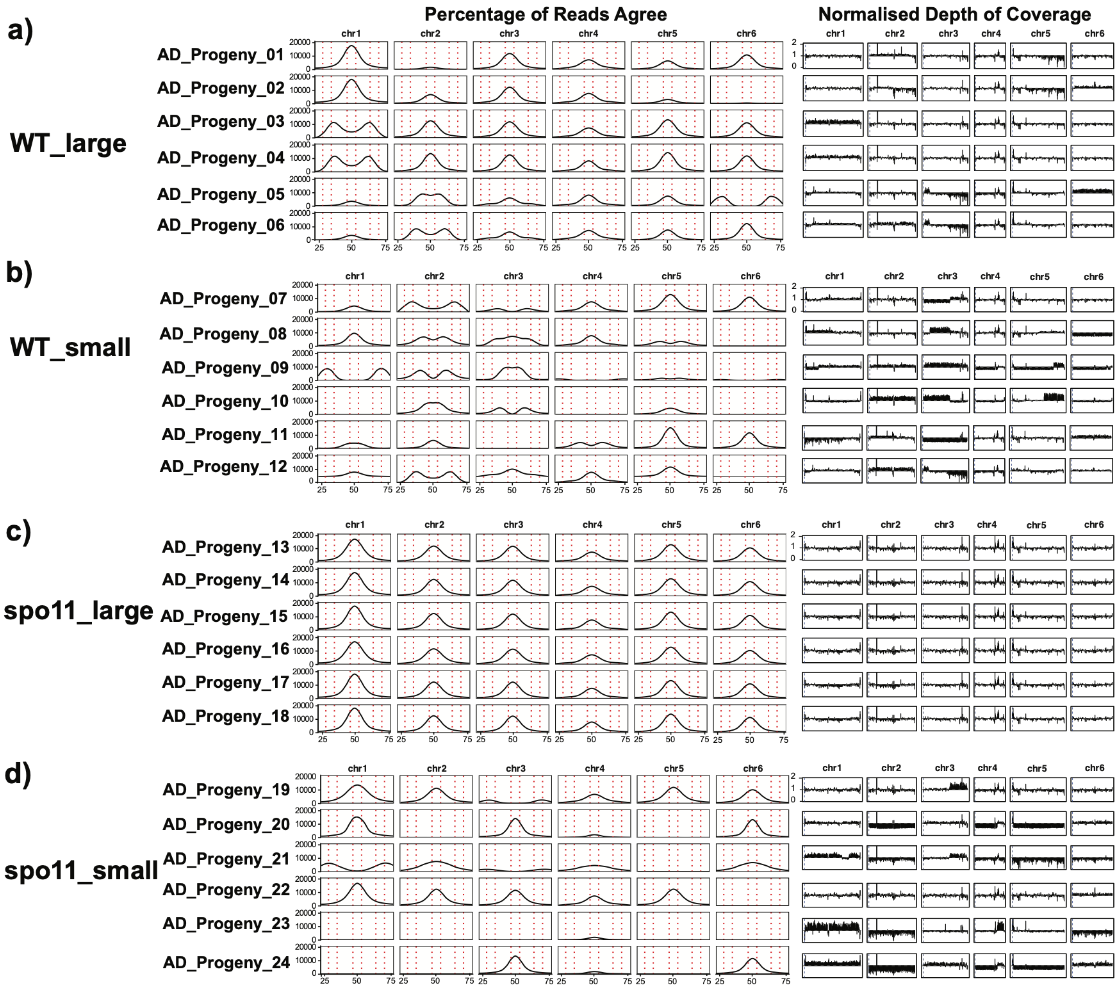


**Figure S11 Aligning *C. neoformans* hybrids crosses to *C. neoformans* H99 reveal copy number and ploidy variations throughout both WT and spo11Δ progeny** Percent of reads agree plotted across chromosomes (**a**) demonstrate near-uniform diploidy in both WT

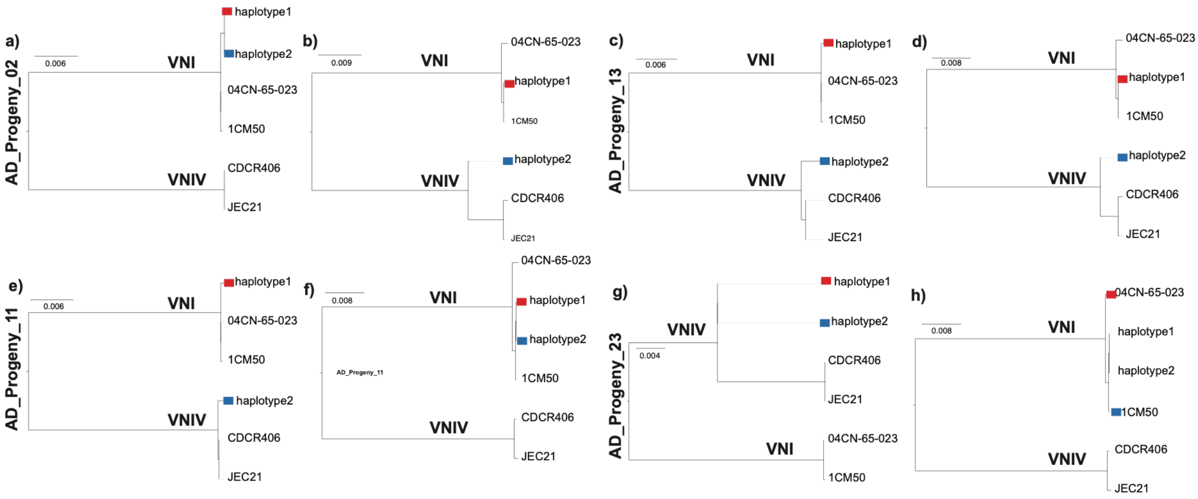


**Figure S12 Phylogenetic analysis of largest haplotypes reveals segregation of haplotypes in both WT and spo11Δ crosses.** Phylogenetic tree of 2 of the largest haplotypes (chromosome 6:1000958-1006704; chromosome 11:486082-492538) of phased hybrid progeny (progeny 2, WT large; progeny 11, WT small; progeny 13, spo11Δ large; progeny 20, spo11Δ small; progeny 23, spo11Δ small)

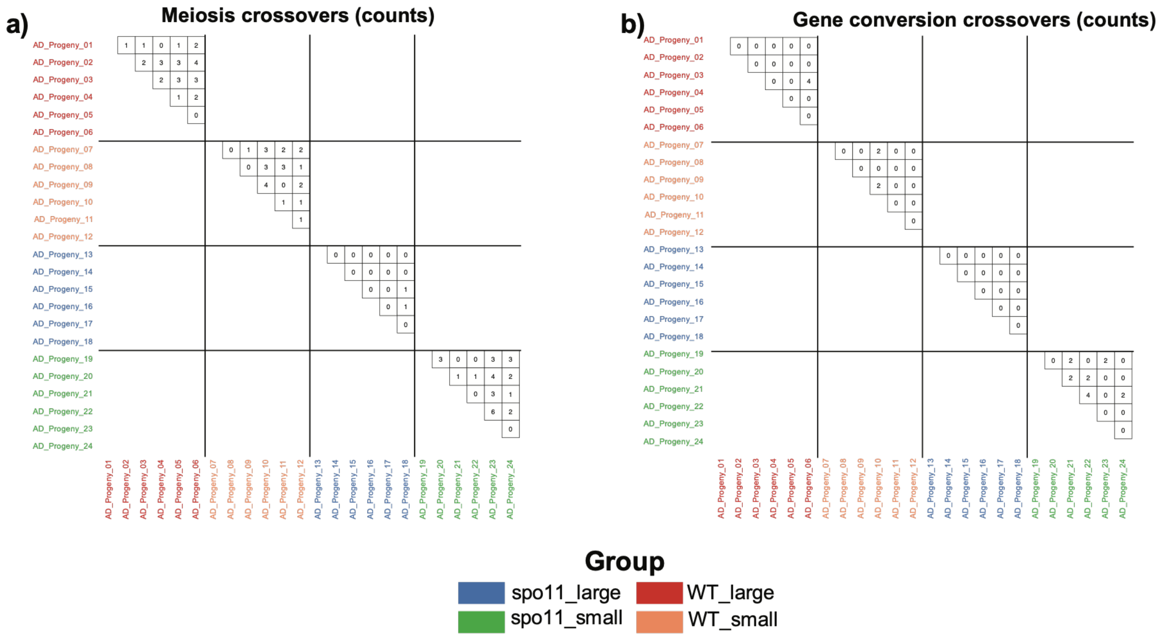


**Figure S13 Counts of crossovers among spo11Δ** progeny. (**a**) We classified crossovers ≥ 1kb from any other crossover and 3 ≥ phased heterozygous positions either side of crossovers as meiosis candidates. (**b**) all remaining crossovers were considered gene conversion crossovers/events.

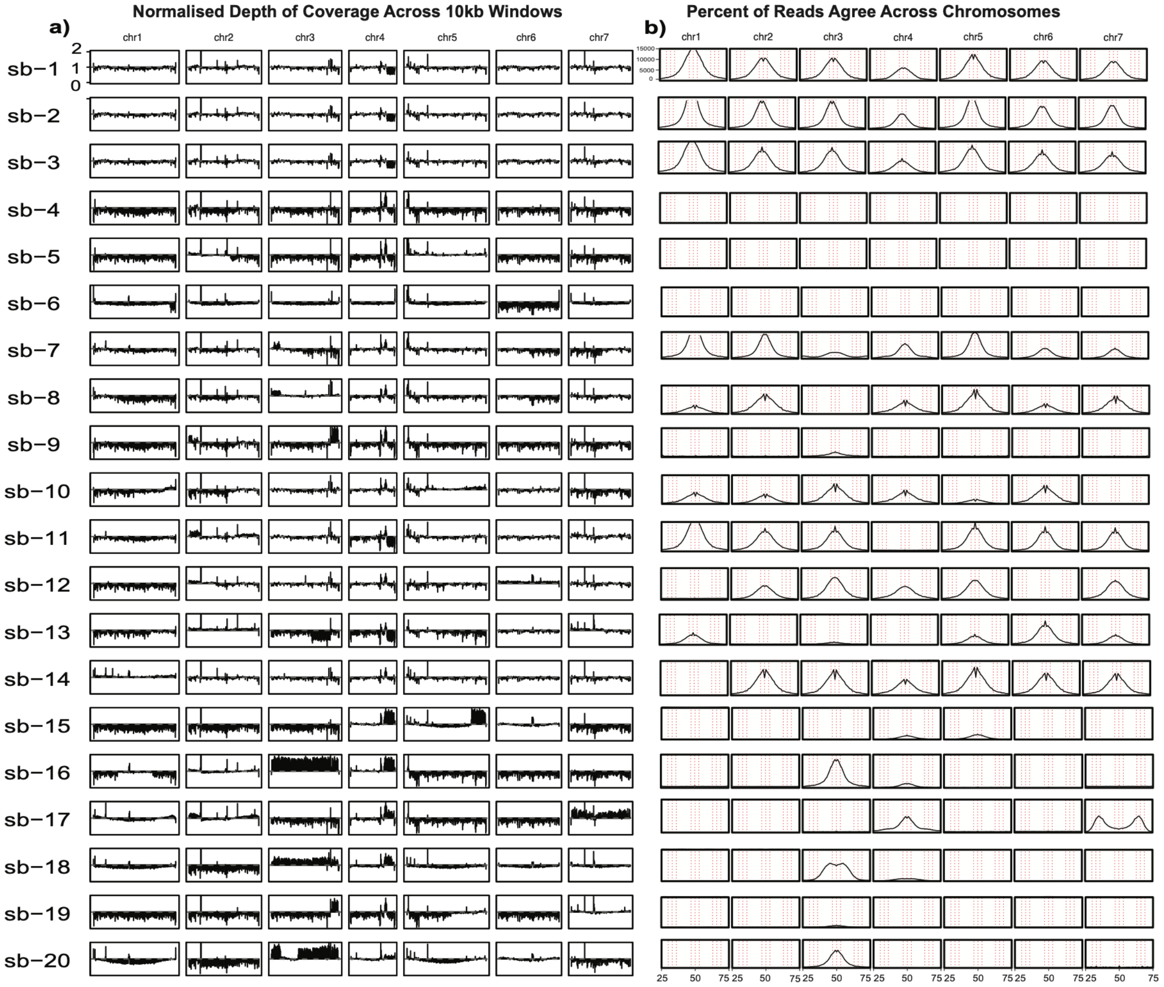


**Figure S14 AD hybrid progeny from Kwon-Chung and Varma (2006) demonstrate ploidy variation and evidence of concerted chromosomal loss.** Alignments *C. neoformans* x *C. deneoformans* progeny to H99 *C. neoformans* reference genome used to plot (**a**) normalised depth of coverage and (**b**) tally of percent of reads agreeing to the reference base.

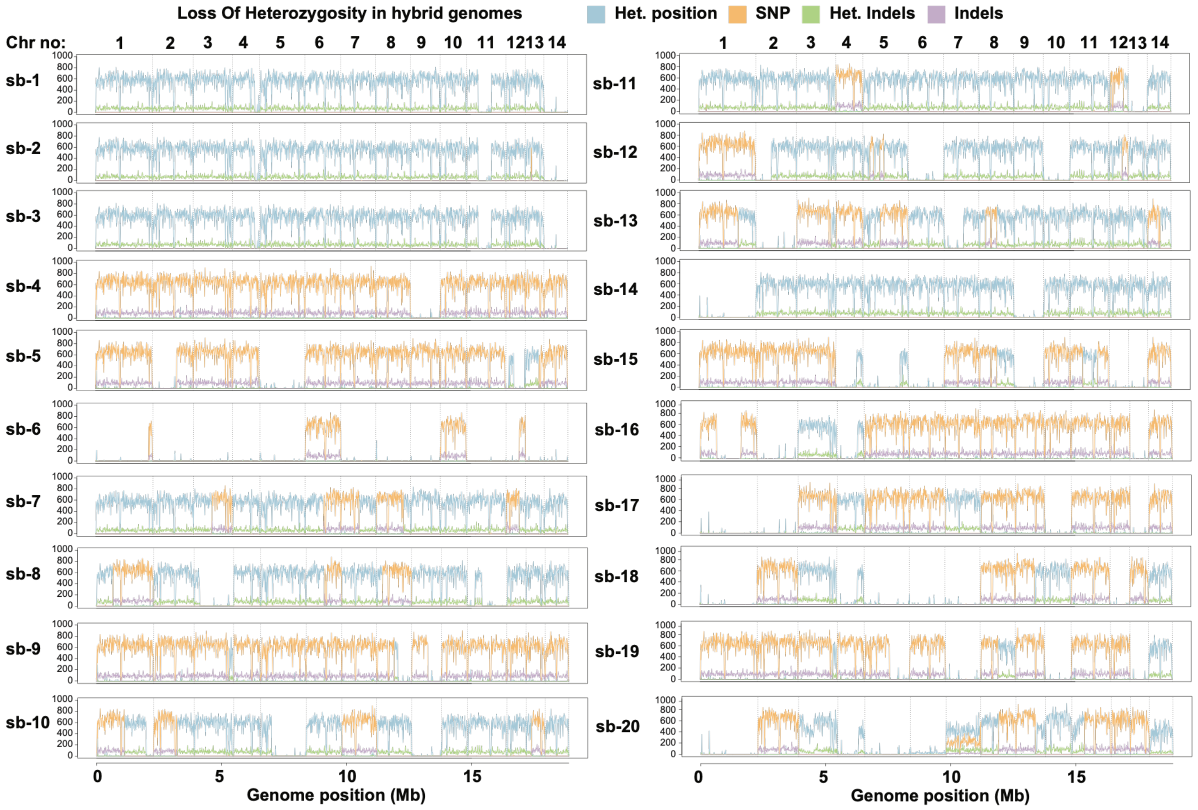


**Figure S15 LOH Observed across aneuploid hybrid progeny.** Variants plotted across the genomes of 20 *C. neoformans* isolates taken from ncbi sra database under the bioproject PRJNA999197 with (blue: heterozygous positions; orange: homozygous SNPs; green: heterozygous indels; purple: homozygous indels). We see large genomics regions where there is a significant drop in heterozygous positions and heterozygous indels, which sometimes are also accompanied by long runs of homozygosity (LROH).

### Supplementary Tables

| Name | GATK ploidy | amb | deletion | het deletion | het insertion | het | insertion | reference | snp | snp multi | BOC (%) |
|---|---|---|---|---|---|---|---|---|---|---|---|
| 04CN-64-029 | Haploid | 474,125 | 2,735 | 0 | 0 | 0 | 2,937 | 17,978,331 | 45,434 | 746 | 95.44 |
| 04CN-64-097 | Haploid | 313,172 | 2,805 | 0 | 0 | 0 | 3,058 | 18,067,747 | 46,359 | 808 | 95.92 |
| 04CN-64-127 | Haploid | 753,188 | 2,788 | 0 | 0 | 0 | 2,922 | 17,674,723 | 46,921 | 767 | 93.85 |
| 04CN-65-023 | Haploid | 297,056 | 2,891 | 0 | 0 | 0 | 3,055 | 18,149,849 | 46,840 | 782 | 96.36 |
| 04CN-65-046 | Haploid | 392,953 | 2,776 | 0 | 0 | 0 | 3,013 | 17,985,939 | 46,266 | 788 | 95.49 |
| 125.91 | Haploid | 382,581 | 2,843 | 0 | 0 | 0 | 3,045 | 17,862,853 | 48,161 | 777 | 94.85 |
| 1CM50 | Haploid | 4,288,901 | 2,623 | 0 | 0 | 0 | 2,730 | 14,009,745 | 46,075 | 715 | 74.44 |
| 881205 | Diploid | 1,053,696 | 9,502 | 115,968 | 20,065 | 951,809 | 9,559 | 15,824,899 | 179,483 | 1,424 | 90.59 |
| A2-102-5 | Haploid | 330,048 | 2,768 | 0 | 0 | 0 | 2,957 | 17,926,975 | 45,945 | 799 | 95.18 |
| ACTA2006-D | Haploid | 238,586 | 15,236 | 0 | 0 | 0 | 16,387 | 17,600,771 | 289,275 | 4,180 | 94.89 |
| ACTA2039-D | Haploid | 242,691 | 15,182 | 0 | 0 | 0 | 16,349 | 17,575,114 | 288,779 | 4,193 | 94.75 |
| ACTA2041-D | Haploid | 221,513 | 15,193 | 0 | 0 | 0 | 16,385 | 17,581,524 | 288,885 | 4,187 | 94.79 |
| ACTA2232-D | Haploid | 215,344 | 15,232 | 0 | 0 | 0 | 16,449 | 17,556,759 | 289,739 | 4,185 | 94.66 |
| ATCC48184 | Diploid | 1,654,378 | 16,931 | 80,206 | 13,777 | 709,725 | 17,176 | 15,186,603 | 303,319 | 1,744 | 86.44 |
| B3501 | Haploid | 1,948,972 | 67,256 | 0 | 0 | 0 | 66,789 | 13,553,833 | 1,065,053 | 3,762 | 78.12 |
| BK158 | Haploid | 335,607 | 2,771 | 0 | 0 | 0 | 2,950 | 18,025,077 | 46,453 | 776 | 95.70 |
| BM08B1 | Haploid | 305,138 | 11,995 | 0 | 0 | 0 | 12,495 | 17,749,294 | 219,524 | 3,245 | 95.27 |
| BM40B3 | Haploid | 267,603 | 11,558 | 0 | 0 | 0 | 13,483 | 17,761,646 | 218,726 | 3,275 | 95.33 |
| Bt100 | Haploid | 540,534 | 2,174 | 0 | 0 | 0 | 2,271 | 17,773,452 | 34,860 | 594 | 94.30 |
| Bt101 | Haploid | 490,940 | 2,079 | 0 | 0 | 0 | 2,172 | 17,825,090 | 32,417 | 517 | 94.56 |
| Bt104 | Haploid | 814,300 | 2,076 | 0 | 0 | 0 | 2,165 | 17,492,396 | 32,419 | 532 | 92.79 |
| Bt106 | Haploid | 593,418 | 2,073 | 0 | 0 | 0 | 2,166 | 17,746,981 | 32,494 | 503 | 94.14 |
| Bt108 | Haploid | 495,109 | 2,157 | 0 | 0 | 0 | 2,232 | 17,798,024 | 34,084 | 555 | 94.42 |
| Bt11 | Haploid | 593,956 | 2,088 | 0 | 0 | 0 | 2,184 | 17,720,586 | 32,414 | 531 | 94.00 |
| Bt110 | Haploid | 551,078 | 2,264 | 0 | 0 | 0 | 2,256 | 17,756,267 | 35,064 | 568 | 94.21 |
| Bt114 | Haploid | 764,900 | 2,424 | 0 | 0 | 0 | 2,443 | 17,552,045 | 37,595 | 626 | 93.14 |
| Bt116 | Haploid | 514,395 | 2,418 | 0 | 0 | 0 | 2,458 | 17,802,110 | 37,579 | 631 | 94.47 |
| Bt117 | Haploid | 595,416 | 2,094 | 0 | 0 | 0 | 2,278 | 17,738,045 | 33,559 | 569 | 94.10 |
| Bt118 | Haploid | 698,839 | 2,072 | 0 | 0 | 0 | 2,162 | 17,631,941 | 32,547 | 510 | 93.53 |
| Bt120 | Haploid | 379,171 | 2,163 | 0 | 0 | 0 | 2,223 | 17,935,249 | 34,093 | 555 | 95.15 |
| Bt121 | Haploid | 815,828 | 2,132 | 0 | 0 | 0 | 2,188 | 17,471,100 | 34,006 | 542 | 92.69 |
| Bt123 | Haploid | 561,694 | 2,195 | 0 | 0 | 0 | 2,269 | 17,747,528 | 34,343 | 565 | 94.16 |
| Bt126 | Haploid | 507,978 | 2,107 | 0 | 0 | 0 | 2,165 | 17,813,561 | 32,737 | 527 | 94.50 |
| Bt129 | Haploid | 617,605 | 2,173 | 0 | 0 | 0 | 2,197 | 17,687,891 | 34,134 | 551 | 93.84 |
| Bt130 | Haploid | 456,831 | 2,267 | 0 | 0 | 0 | 2,418 | 17,805,420 | 37,424 | 601 | 94.48 |
| Bt137 | Haploid | 650,201 | 2,116 | 0 | 0 | 0 | 2,220 | 17,658,292 | 33,345 | 544 | 93.68 |
| Bt15 | Haploid | 348,939 | 2,091 | 0 | 0 | 0 | 2,204 | 17,993,157 | 32,716 | 523 | 95.45 |
| Bt150 | Haploid | 420,581 | 2,166 | 0 | 0 | 0 | 2,253 | 17,906,218 | 34,015 | 563 | 94.99 |
| Bt151 | Haploid | 650,538 | 2,117 | 0 | 0 | 0 | 2,207 | 17,663,594 | 33,315 | 546 | 93.71 |
| Bt153 | Haploid | 830,803 | 2,132 | 0 | 0 | 0 | 2,204 | 17,465,131 | 34,127 | 547 | 92.66 |
| Bt155 | Haploid | 468,810 | 2,133 | 0 | 0 | 0 | 2,298 | 17,879,744 | 33,753 | 570 | 94.85 |
| Bt156 | Haploid | 497,169 | 2,388 | 0 | 0 | 0 | 2,591 | 17,795,265 | 39,572 | 657 | 94.44 |
| Bt160 | Haploid | 577,630 | 2,094 | 0 | 0 | 0 | 2,182 | 17,727,549 | 32,526 | 538 | 94.04 |
| Bt161 | Haploid | 612,558 | 2,241 | 0 | 0 | 0 | 2,315 | 17,690,726 | 35,448 | 588 | 93.86 |
| Bt17 | Haploid | 384,325 | 2,104 | 0 | 0 | 0 | 2,221 | 17,933,568 | 32,571 | 526 | 95.13 |
| Bt19 | Haploid | 600,836 | 2,091 | 0 | 0 | 0 | 2,179 | 17,684,856 | 32,442 | 533 | 93.81 |
| Bt19_v2 | Haploid | 554,491 | 2,084 | 0 | 0 | 0 | 2,193 | 17,787,612 | 33,746 | 566 | 94.36 |
| Bt2 | Haploid | 729,372 | 2,251 | 0 | 0 | 0 | 2,333 | 17,585,201 | 35,315 | 588 | 93.30 |
| Bt20 | Haploid | 684,741 | 2,257 | 0 | 0 | 0 | 2,323 | 17,666,396 | 36,042 | 602 | 93.74 |
| Bt202 | Haploid | 515,734 | 2,094 | 0 | 0 | 0 | 2,188 | 17,797,929 | 32,414 | 531 | 94.41 |
| Bt203 | Haploid | 638,493 | 2,153 | 0 | 0 | 0 | 2,193 | 17,650,985 | 34,107 | 550 | 93.64 |
| Bt205 | Haploid | 431,917 | 2,093 | 0 | 0 | 0 | 2,197 | 17,872,956 | 32,432 | 528 | 94.81 |
| Bt207 | Haploid | 476,656 | 2,170 | 0 | 0 | 0 | 2,223 | 17,843,606 | 34,034 | 564 | 94.66 |
| Bt209 | Haploid | 760,483 | 2,074 | 0 | 0 | 0 | 2,162 | 17,526,594 | 32,448 | 528 | 92.98 |
| Bt210 | Haploid | 335,338 | 2,120 | 0 | 0 | 0 | 2,225 | 18,023,801 | 32,855 | 550 | 95.61 |
| Bt211 | Haploid | 674,392 | 2,159 | 0 | 0 | 0 | 2,213 | 17,620,801 | 34,067 | 548 | 93.48 |
| Bt213 | Haploid | 1,072,543 | 2,147 | 0 | 0 | 0 | 2,232 | 17,258,808 | 33,896 | 553 | 91.57 |

| | | | | | | | | | | |
|---|---|---|---|---|---|---|---|---|---|---|
| Bt23 | Haploid | 721,531 | 2,109 | 0 | 0 | 0 | 2,247 | 17,571,812 | 33,569 | 564 | 93.22 |
| Bt30 | Haploid | 479,253 | 2,258 | 0 | 0 | 0 | 2,314 | 17,777,044 | 35,775 | 603 | 94.32 |
| Bt37 | Haploid | 547,961 | 2,189 | 0 | 0 | 0 | 2,280 | 17,778,950 | 34,768 | 556 | 94.33 |
| Bt42 | Haploid | 603,418 | 2,085 | 0 | 0 | 0 | 2,161 | 17,717,158 | 32,352 | 518 | 93.98 |
| Bt43 | Haploid | 506,698 | 2,175 | 0 | 0 | 0 | 2,256 | 17,840,480 | 34,169 | 562 | 94.65 |
| Bt44 | Haploid | 625,976 | 2,256 | 0 | 0 | 0 | 2,243 | 17,680,267 | 35,062 | 567 | 93.80 |
| Bt47 | Haploid | 483,242 | 2,079 | 0 | 0 | 0 | 2,172 | 17,857,934 | 32,608 | 517 | 94.73 |
| Bt57 | Haploid | 422,758 | 2,113 | 0 | 0 | 0 | 2,177 | 17,887,563 | 32,746 | 528 | 94.89 |
| Bt59 | Haploid | 885,484 | 2,061 | 0 | 0 | 0 | 2,161 | 17,418,896 | 32,373 | 524 | 92.41 |
| Bt62 | Haploid | 394,418 | 2,138 | 0 | 0 | 0 | 2,202 | 17,926,846 | 32,788 | 522 | 95.10 |
| Bt68 | Haploid | 537,086 | 2,166 | 0 | 0 | 0 | 2,209 | 17,755,202 | 34,140 | 546 | 94.20 |
| Bt72 | Haploid | 713,303 | 2,107 | 0 | 0 | 0 | 2,286 | 17,628,093 | 33,538 | 568 | 93.52 |
| Bt77 | Haploid | 567,435 | 2,121 | 0 | 0 | 0 | 2,274 | 17,760,574 | 33,655 | 567 | 94.22 |
| Bt78 | Haploid | 689,613 | 2,074 | 0 | 0 | 0 | 2,156 | 17,600,180 | 32,328 | 522 | 93.36 |
| Bt79 | Haploid | 805,373 | 2,063 | 0 | 0 | 0 | 2,175 | 17,497,573 | 32,358 | 530 | 92.82 |
| Bt8 | Haploid | 725,457 | 2,079 | 0 | 0 | 0 | 2,164 | 17,605,823 | 32,531 | 503 | 93.40 |
| Bt83 | Haploid | 935,760 | 2,101 | 0 | 0 | 0 | 2,235 | 17,362,662 | 33,415 | 566 | 92.11 |
| Bt87 | Haploid | 574,717 | 2,073 | 0 | 0 | 0 | 2,184 | 17,731,853 | 32,417 | 533 | 94.06 |
| Bt90 | Haploid | 793,896 | 2,105 | 0 | 0 | 0 | 2,161 | 17,516,452 | 32,605 | 518 | 92.92 |
| Bt93 | Haploid | 1,004,208 | 2,228 | 0 | 0 | 0 | 2,199 | 17,257,350 | 34,351 | 570 | 91.56 |
| Bt95 | Haploid | 481,899 | 2,103 | 0 | 0 | 0 | 2,281 | 17,840,012 | 33,603 | 573 | 94.64 |
| Bt97 | Haploid | 572,655 | 2,111 | 0 | 0 | 0 | 2,261 | 17,764,229 | 33,559 | 569 | 94.24 |
| CB-100-1 | Diploid | 401,787 | 7,716 | 111,482 | 18,419 | 938,136 | 7,724 | 16,999,663 | 107,163 | 418 | 96.29 |
| CB-100-2 | Diploid | 564,453 | 12,580 | 108,999 | 18,083 | 892,131 | 12,808 | 16,674,846 | 181,924 | 648 | 94.77 |
| CB-100-3 | Diploid | 540,092 | 7,375 | 110,501 | 18,207 | 925,273 | 7,290 | 16,832,423 | 105,536 | 399 | 95.32 |
| CB-100-5 | Diploid | 461,942 | 10,322 | 109,901 | 18,309 | 909,913 | 10,334 | 16,848,555 | 148,198 | 523 | 95.58 |
| CB-100-6 | Diploid | 527,804 | 12,605 | 108,582 | 17,832 | 898,646 | 12,663 | 16,711,132 | 171,886 | 659 | 94.94 |
| CBS132 | Diploid | 1,398,543 | 24,647 | 64,607 | 11,154 | 558,791 | 24,669 | 15,338,673 | 378,989 | 1,457 | 86.83 |
| CBS6995 | Haploid | 2,521,385 | 60,536 | 0 | 0 | 0 | 61,382 | 13,138,182 | 1,023,072 | 3,423 | 75.63 |
| CBS7748 | Haploid | 1,543,200 | 73,371 | 0 | 0 | 0 | 73,347 | 14,033,512 | 1,099,462 | 4,065 | 80.91 |
| CBS7821 | Diploid | 1,449,437 | 421 | 109,862 | 18,137 | 979,907 | 443 | 15,958,876 | 8,705 | 76 | 90.40 |
| CC-100-1 | Diploid | 403,694 | 7,807 | 113,650 | 18,651 | 940,610 | 7,793 | 16,983,557 | 107,260 | 408 | 96.24 |
| CC-100-2 | Diploid | 498,078 | 12,622 | 106,615 | 17,475 | 887,497 | 12,655 | 16,763,802 | 172,047 | 657 | 95.14 |
| CC-100-3 | Diploid | 493,170 | 18,010 | 89,723 | 14,981 | 744,673 | 18,087 | 16,787,008 | 247,867 | 942 | 94.87 |
| CC-100-4 | Diploid | 517,333 | 15,786 | 93,451 | 15,490 | 780,699 | 15,758 | 16,732,378 | 213,782 | 843 | 94.59 |
| CC-100-6 | Diploid | 666,361 | 11,685 | 97,218 | 15,949 | 818,425 | 11,607 | 16,634,173 | 167,649 | 626 | 94.00 |
| CCTP15 | Haploid | 290,204 | 12,344 | 0 | 0 | 0 | 12,649 | 17,797,938 | 223,478 | 3,304 | 95.55 |
| CCTP44 | Diploid | 522,143 | 6,997 | 133,769 | 23,495 | 1,072,035 | 7,097 | 16,487,183 | 134,150 | 1,405 | 94.58 |
| CCTP50 | Diploid | 573,982 | 6,798 | 132,099 | 22,959 | 1,072,738 | 6,917 | 16,381,393 | 131,855 | 1,357 | 93.99 |
| CDCR406 | Haploid | 2,147,798 | 66,439 | 0 | 0 | 0 | 66,034 | 13,385,868 | 1,054,310 | 3,742 | 77.16 |
| CN-100-1 | Diploid | 464,693 | 7,679 | 113,502 | 18,758 | 946,396 | 7,674 | 16,898,369 | 107,331 | 413 | 95.81 |
| CN-100-2 | Diploid | 559,950 | 9,100 | 104,636 | 17,386 | 860,325 | 9,151 | 16,871,375 | 132,263 | 474 | 95.31 |
| CN-100-3 | Diploid | 429,895 | 7,884 | 115,396 | 19,105 | 955,098 | 7,845 | 16,931,518 | 107,734 | 428 | 96.05 |
| CN-100-4 | Diploid | 440,190 | 7,737 | 112,929 | 18,655 | 941,069 | 7,713 | 16,931,545 | 107,044 | 414 | 95.96 |
| CN-100-5 | Diploid | 458,614 | 7,552 | 114,452 | 18,874 | 948,322 | 7,557 | 16,893,964 | 107,015 | 412 | 95.80 |
| CN-100-6 | Diploid | 420,086 | 7,683 | 116,124 | 19,150 | 955,017 | 7,688 | 16,959,433 | 106,957 | 417 | 96.20 |
| CW-100-1 | Diploid | 423,391 | 8,042 | 104,236 | 17,224 | 868,513 | 7,959 | 16,996,969 | 110,552 | 427 | 95.89 |
| CW-100-2 | Diploid | 541,844 | 7,624 | 115,468 | 18,922 | 954,030 | 7,563 | 16,814,452 | 106,883 | 407 | 95.42 |
| CW-100-3 | Diploid | 558,934 | 12,437 | 100,236 | 16,502 | 827,907 | 12,495 | 16,756,771 | 171,406 | 669 | 94.75 |
| CW-100-4 | Diploid | 483,268 | 7,806 | 120,919 | 19,657 | 967,585 | 7,765 | 16,868,158 | 108,418 | 411 | 95.82 |
| CW-100-5 | Diploid | 542,993 | 7,603 | 117,058 | 19,132 | 962,312 | 7,588 | 16,783,712 | 106,776 | 400 | 95.31 |
| CW-100-6 | Diploid | 389,877 | 1,634 | 115,333 | 18,912 | 958,860 | 1,726 | 17,158,849 | 23,132 | 116 | 96.76 |
| CY-100-1 | Diploid | 472,715 | 7,603 | 107,933 | 17,889 | 906,160 | 7,797 | 16,887,891 | 107,799 | 402 | 95.51 |
| CY-100-2 | Diploid | 485,465 | 11,428 | 110,279 | 18,232 | 910,604 | 11,582 | 16,745,384 | 153,961 | 582 | 95.08 |
| CY-100-3 | Diploid | 368,781 | 1,736 | 122,377 | 20,066 | 1,004,236 | 1,826 | 17,091,368 | 23,444 | 125 | 96.69 |
| CY-100-4 | Diploid | 498,092 | 11,403 | 97,922 | 16,337 | 823,638 | 11,606 | 16,807,586 | 158,651 | 602 | 94.90 |
| CY-100-5 | Diploid | 464,354 | 12,287 | 98,988 | 16,483 | 820,578 | 12,227 | 16,879,212 | 169,075 | 676 | 95.34 |
| CY-100-6 | Diploid | 396,233 | 1,776 | 127,353 | 20,719 | 1,040,012 | 1,873 | 17,069,230 | 23,429 | 124 | 96.79 |
| D17-1 | Haploid | 247,908 | 2,766 | 0 | 0 | 0 | 2,971 | 18,043,613 | 45,842 | 804 | 95.79 |
| EM3 | Haploid | 1,024,871 | 82,773 | 0 | 0 | 0 | 83,246 | 14,676,375 | 1,139,177 | 4,335 | 84.62 |

| | | | | | | | | | | | |
|---|---|---|---|---|---|---|---|---|---|---|---|
| FN62 | Diploid | 491,811 | 12,226 | 109,393 | 19,410 | 876,416 | 12,323 | 16,600,139 | 203,681 | 1,782 | 94.41 |
| Gb159-1 | Haploid | 557,970 | 2,154 | 0 | 0 | 0 | 2,271 | 17,767,197 | 34,261 | 601 | 94.26 |
| HAMDAN | Haploid | 3,144,129 | 10,507 | 0 | 0 | 0 | 11,074 | 14,855,886 | 214,575 | 2,912 | 79.91 |
| IFN-R21 | Diploid | 551,557 | 13,824 | 125,280 | 22,116 | 987,386 | 13,850 | 16,328,880 | 226,169 | 1,613 | 93.80 |
| IFN-R23 | Diploid | 532,362 | 11,203 | 130,516 | 22,944 | 1,029,006 | 11,243 | 16,394,080 | 188,354 | 1,530 | 94.17 |
| IFN-R26 | Diploid | 524,832 | 16,052 | 106,746 | 18,850 | 830,856 | 15,943 | 16,388,436 | 256,444 | 1,846 | 93.35 |
| IFN26 | Diploid | 640,917 | 6,761 | 130,166 | 22,618 | 1,067,708 | 6,843 | 16,292,419 | 130,593 | 1,334 | 93.48 |
| IFN82 | Haploid | 301,857 | 11,945 | 0 | 0 | 0 | 12,338 | 17,767,835 | 218,197 | 3,251 | 95.36 |
| JEC20a | Haploid | 698,746 | 87,042 | 0 | 0 | 0 | 88,235 | 15,229,283 | 1,150,554 | 4,512 | 87.66 |
| JEC20a_msh2-1 | Haploid | 705,278 | 86,990 | 0 | 0 | 0 | 88,391 | 15,224,956 | 1,150,594 | 4,541 | 87.64 |
| JEC21 | Haploid | 2,152,223 | 63,161 | 0 | 0 | 0 | 63,606 | 13,501,441 | 1,042,143 | 3,471 | 77.68 |
| KW5 | Haploid | 1,243,897 | 11,410 | 0 | 0 | 0 | 12,050 | 16,769,126 | 223,244 | 3,133 | 90.09 |
| LP-RSA116 | Haploid | 683,503 | 2,242 | 0 | 0 | 0 | 2,295 | 17,645,041 | 36,518 | 612 | 93.63 |
| LP-RSA1343 | Haploid | 607,662 | 2,261 | 0 | 0 | 0 | 2,321 | 17,710,009 | 35,257 | 578 | 93.96 |
| LP-RSA2296 | Haploid | 642,981 | 2,759 | 0 | 0 | 0 | 2,976 | 17,710,388 | 47,072 | 773 | 94.04 |
| LP-RSA3042 | Haploid | 583,974 | 2,231 | 0 | 0 | 0 | 2,272 | 17,708,089 | 35,923 | 591 | 93.96 |
| LP-RSA848 | Haploid | 460,846 | 2,183 | 0 | 0 | 0 | 2,250 | 17,868,658 | 34,041 | 560 | 94.80 |
| Lu41-1 | Haploid | 284,473 | 11,290 | 0 | 0 | 0 | 13,449 | 17,824,914 | 216,718 | 3,239 | 95.65 |
| Lu42-3 | Haploid | 272,367 | 11,466 | 0 | 0 | 0 | 13,583 | 17,776,671 | 218,715 | 3,247 | 95.41 |
| Lu45-3 | Haploid | 257,250 | 11,303 | 0 | 0 | 0 | 13,419 | 17,828,195 | 216,794 | 3,236 | 95.67 |
| MW-RSA3834 | Haploid | 237,109 | 2,191 | 0 | 0 | 0 | 2,286 | 18,124,750 | 34,023 | 579 | 96.15 |
| MW-RSA524 | Haploid | 262,557 | 2,192 | 0 | 0 | 0 | 2,287 | 18,093,808 | 33,995 | 578 | 95.99 |
| NL14-1 | Haploid | 265,400 | 11,466 | 0 | 0 | 0 | 13,575 | 17,802,585 | 218,989 | 3,260 | 95.55 |
| NL38-1 | Haploid | 264,785 | 11,418 | 0 | 0 | 0 | 13,484 | 17,790,553 | 217,804 | 3,290 | 95.48 |
| O18-1 | Haploid | 334,901 | 11,327 | 0 | 0 | 0 | 13,354 | 17,708,294 | 216,105 | 3,223 | 95.03 |
| O22-1 | Haploid | 253,423 | 11,311 | 0 | 0 | 0 | 13,333 | 17,818,452 | 215,782 | 3,247 | 95.61 |
| P1W10 | Haploid | 1,643,004 | 13,650 | 0 | 0 | 0 | 13,631 | 16,641,709 | 269,609 | 912 | 89.67 |
| P2W10 | Haploid | 1,814,614 | 18,854 | 0 | 0 | 0 | 18,923 | 16,404,478 | 320,173 | 1,146 | 88.74 |
| P31-1 | Haploid | 252,136 | 11,299 | 0 | 0 | 0 | 13,279 | 17,788,937 | 214,374 | 3,174 | 95.45 |
| P3W10 | Haploid | 1,855,144 | 12,871 | 0 | 0 | 0 | 12,692 | 16,522,704 | 260,577 | 926 | 88.98 |
| P43-3 | Haploid | 295,796 | 12,110 | 0 | 0 | 0 | 12,918 | 17,795,226 | 225,738 | 3,338 | 95.55 |
| P4W10 | Haploid | 1,566,092 | 14,079 | 0 | 0 | 0 | 13,852 | 16,702,301 | 286,977 | 983 | 90.09 |
| P5W10 | Haploid | 2,010,982 | 15,225 | 0 | 0 | 0 | 15,123 | 16,215,481 | 284,777 | 1,067 | 87.51 |
| P6W10 | Haploid | 2,787,430 | 12,190 | 0 | 0 | 0 | 11,973 | 15,544,769 | 281,490 | 891 | 83.91 |
| RCT14 | Diploid | 505,323 | 11,559 | 109,298 | 19,200 | 879,961 | 11,878 | 16,548,115 | 198,953 | 1,785 | 94.12 |
| RCT22 | Haploid | 301,661 | 12,110 | 0 | 0 | 0 | 12,544 | 17,776,880 | 220,604 | 3,274 | 95.42 |
| RCT26 | Haploid | 266,361 | 11,944 | 0 | 0 | 0 | 12,440 | 17,884,554 | 218,487 | 3,241 | 95.98 |
| SA8961 | Haploid | 287,962 | 11,994 | 0 | 0 | 0 | 12,484 | 17,759,024 | 219,537 | 3,246 | 95.32 |
| SA8963 | Haploid | 338,415 | 11,940 | 0 | 0 | 0 | 12,411 | 17,799,441 | 218,313 | 3,224 | 95.52 |
| SSD719-vitro | Haploid | 1,379,602 | 18,642 | 0 | 0 | 0 | 18,949 | 16,818,541 | 399,210 | 1,278 | 91.35 |
| SSD719-vivo | Haploid | 2,493,569 | 10,880 | 0 | 0 | 0 | 10,905 | 15,932,553 | 224,971 | 806 | 85.65 |
| Tu241-1 | Haploid | 561,583 | 2,437 | 0 | 0 | 0 | 2,411 | 17,706,937 | 39,282 | 658 | 93.97 |
| Tu259-1 | Haploid | 269,552 | 2,787 | 0 | 0 | 0 | 2,919 | 18,076,803 | 44,506 | 746 | 95.96 |
| UON11536 | Haploid | 870,567 | 14,365 | 0 | 0 | 0 | 15,158 | 16,856,934 | 283,530 | 3,942 | 90.91 |
| V18-2 | Haploid | 386,849 | 11,558 | 0 | 0 | 0 | 13,511 | 17,707,103 | 220,052 | 3,272 | 95.05 |
| V28-1 | Haploid | 310,985 | 11,654 | 0 | 0 | 0 | 13,240 | 17,770,846 | 220,798 | 3,250 | 95.39 |
| V31B1 | Haploid | 307,860 | 12,245 | 0 | 0 | 0 | 12,795 | 17,748,785 | 223,513 | 3,323 | 95.29 |
| V53-4 | Haploid | 330,853 | 12,035 | 0 | 0 | 0 | 12,507 | 17,753,487 | 220,235 | 3,342 | 95.29 |
| VS24A4 | Haploid | 341,515 | 11,669 | 0 | 0 | 0 | 13,179 | 17,715,608 | 220,772 | 3,252 | 95.10 |
| VS32-1 | Haploid | 315,687 | 12,035 | 0 | 0 | 0 | 12,454 | 17,752,906 | 221,046 | 3,241 | 95.29 |
| YX1 | Diploid | 357,986 | 34,304 | 9,263 | 1,874 | 79,684 | 34,359 | 17,196,627 | 461,463 | 1,725 | 94.33 |
| YX10 | Diploid | 373,154 | 14,749 | 107,657 | 17,681 | 878,481 | 14,734 | 16,843,470 | 196,866 | 773 | 95.68 |
| YX12 | Diploid | 303,613 | 15,712 | 69,522 | 11,784 | 581,717 | 15,888 | 17,232,329 | 207,089 | 821 | 96.00 |
| YX14 | Diploid | 339,551 | 7,054 | 107,207 | 17,769 | 905,437 | 7,360 | 17,010,793 | 94,013 | 332 | 96.08 |
| YX16 | Diploid | 313,591 | 16,870 | 64,653 | 11,043 | 543,877 | 16,887 | 17,214,864 | 221,936 | 868 | 95.77 |
| YX17 | Diploid | 259,507 | 785 | 103,655 | 17,343 | 866,879 | 831 | 17,370,548 | 9,962 | 51 | 97.24 |
| YX19 | Diploid | 277,823 | 5,137 | 98,535 | 16,525 | 829,709 | 5,150 | 17,295,268 | 70,615 | 224 | 96.98 |
| YX3 | Diploid | 259,803 | 13,509 | 27,781 | 4,998 | 249,679 | 14,018 | 17,611,038 | 182,476 | 664 | 95.84 |
| YX4 | Diploid | 270,048 | 7,592 | 87,112 | 14,677 | 740,853 | 7,630 | 17,366,881 | 100,968 | 380 | 97.01 |
| YX43 | Diploid | 312,823 | 8,728 | 103,191 | 17,058 | 868,478 | 8,763 | 17,113,257 | 117,048 | 431 | 96.54 |
| YX44 | Diploid | 285,096 | 245 | 124,244 | 20,574 | 1,040,496 | 179 | 17,212,261 | 3,124 | 25 | 97.41 |
| YX45 | Diploid | 302,973 | 4,577 | 113,097 | 18,907 | 951,830 | 4,768 | 17,135,171 | 63,534 | 237 | 96.83 |
| YX47 | Diploid | 141,564 | 295 | 74,554 | 11,203 | 905,047 | 224 | 17,296,255 | 3,437 | 48 | 96.83 |
| YX48 | Diploid | 302,647 | 4,634 | 112,924 | 18,770 | 952,138 | 4,780 | 17,138,755 | 63,658 | 231 | 96.85 |
| YX49 | Diploid | 330,930 | 10,980 | 109,338 | 18,258 | 913,232 | 11,044 | 17,014,047 | 145,305 | 580 | 96.46 |
| YX5 | Diploid | 265,188 | 7,591 | 85,861 | 14,406 | 721,614 | 7,601 | 17,388,691 | 100,945 | 378 | 97.02 |
| YX56 | Diploid | 283,609 | 3,582 | 105,005 | 17,383 | 889,609 | 3,533 | 17,265,102 | 43,976 | 237 | 97.02 |
| YX58 | Diploid | 277,975 | 236 | 119,056 | 19,790 | 990,120 | 153 | 17,280,580 | 2,688 | 21 | 97.47 |
| YX59 | Diploid | 285,232 | 227 | 124,111 | 20,568 | 1,039,892 | 160 | 17,218,925 | 2,700 | 24 | 97.44 |
| YX6 | Diploid | 433,745 | 41,237 | 36,098 | 6,396 | 298,055 | 41,588 | 16,556,289 | 554,207 | 2,050 | 92.83 |
| YX60 | Diploid | 270,303 | 243 | 116,840 | 19,307 | 969,174 | 160 | 17,313,480 | 2,737 | 21 | 97.52 |
| YX61 | Diploid | 337,802 | 14,517 | 98,105 | 16,458 | 817,513 | 14,438 | 16,987,188 | 197,926 | 692 | 96.06 |
| YX64 | Diploid | 322,577 | 3,491 | 102,957 | 16,986 | 865,332 | 3,432 | 17,288,880 | 43,831 | 232 | 97.01 |
| YX65 | Diploid | 292,228 | 219 | 123,296 | 20,387 | 1,035,431 | 155 | 17,213,743 | 2,737 | 25 | 97.38 |
| YX7 | Diploid | 273,791 | 7,550 | 88,483 | 14,828 | 744,750 | 7,571 | 17,355,758 | 101,011 | 388 | 96.98 |
| YX70 | Diploid | 313,844 | 3,502 | 103,929 | 17,162 | 878,302 | 3,466 | 17,290,266 | 43,673 | 227 | 97.09 |
| ZC0004 | Haploid | 304,385 | 12,186 | 0 | 0 | 0 | 12,980 | 17,820,831 | 227,344 | 3,395 | 95.69 |
| ZC0007 | Haploid | 304,517 | 12,009 | 0 | 0 | 0 | 12,555 | 17,819,129 | 221,323 | 3,252 | 95.65 |
| ZC0015 | Haploid | 307,912 | 12,175 | 0 | 0 | 0 | 12,568 | 17,750,582 | 222,719 | 3,309 | 95.29 |
| ZC0021 | Haploid | 323,586 | 11,881 | 0 | 0 | 0 | 12,788 | 17,786,074 | 224,504 | 3,318 | 95.49 |

**Table S1.** Details of variant calling of 197 isolates analysed in this study using GATK (amb = ambiguous, het = heterozygous, snp multi = multiple (often consecutive) SNPs reported as a single VCF entry, BOC = Breadth of Coverage).

| Isolate | Sequence | Biosample Accession | Bioproject Accession | Length (Mb) |
|---|---|---|---|---|
| 881205 | Hap1 | SAMN51237234 | PRJNA1352182 | 18.94 |
| | Hap2 | | PRJNA1353001 | 18.91 |
| | Mito | - | - | 0.03 |
| ATCC48184 | Hap1 | SAMN51237237 | PRJNA1350146 | 20.42 |
| | Hap2 | | PRJNA1350155 | 22.07 |
| | Mito | - | - | 0.03 |
| CBS132 | Hap1 | SAMN51237236 | PRJNA1350895 | 19.22 |
| | Hap2 | | PRJNA1350980 | 17.83 |
| | Mito | - | - | 0.03 |
| CBS7821 | Hap1 | SAMN51237235 | PRJNA1351368 | 19.29 |
| | Hap2 | | PRJNA1351898 | 20.74 |
| | Mito | - | - | 0.03 |

**Table S2.** Variable lengths of *C. neoformans* long-read assemblies. Hybrid assemblies consisting of haplotype 1, haplotype 2 and mitochondrial sequences were assembled.

| Isolate | < 25 % | < 50 % | < 75 % | > 75 % | No match |
|---|---|---|---|---|---|
| 881205 | 1.08 | 83.68 | 0.13 | 8.98 | 6.13 |
| ATCC58184 | 0.80 | 41.03 | 0.41 | 45.92 | 11.84 |
| CBS132 | 1.04 | 42.73 | 0.85 | 33.45 | 21.93 |
| CBS7821 | 5.92 | 75.42 | 0.29 | 6.11 | 12.26 |

**Table S3.** Alignment percentages of haplotypes when compared to each other of hybrid phased assemblies.

| Isolate A | Isolate B | OPG | OPP | OPP Same | OPP Cross | OPP Cross (GC) | OPP Cross (Meiosis) | OPP Cross (%) | OPP Cross (GC)% | OPP Cross (Meiosis)% |
|---|---|---|---|---|---|---|---|---|---|---|
| D018 | D123 | 1697 | 6623 | 6593 | 27 | 2 | 5 | 0.41 | 7.41 | 18.52 |
| D018 | D124 | 1700 | 6627 | 6597 | 27 | 2 | 5 | 0.41 | 7.41 | 18.52 |
| D018 | D125 | 1674 | 6578 | 6556 | 19 | 0 | 6 | 0.29 | 0 | 31.58 |
| D018 | D127 | 1578 | 6096 | 6067 | 26 | 4 | 4 | 0.43 | 15.38 | 15.38 |
| D018 | D129 | 1603 | 6164 | 6147 | 15 | 2 | 0 | 0.24 | 13.33 | 0 |
| D018 | D131 | 1589 | 6168 | 6149 | 16 | 2 | 1 | 0.26 | 12.5 | 6.25 |
| D018 | D132 | 1611 | 6256 | 6235 | 18 | 2 | 4 | 0.29 | 11.11 | 22.22 |
| D018 | D133 | 1686 | 6582 | 6563 | 16 | 0 | 2 | 0.24 | 0 | 12.5 |
| D018 | D134 | 1820 | 7040 | 7024 | 14 | 0 | 2 | 0.2 | 0 | 14.29 |
| D123 | D018 | 1697 | 6623 | 6593 | 27 | 2 | 7 | 0.41 | 7.41 | 25.93 |
| D123 | D124 | 3057 | 12266 | 12266 | 0 | 0 | 0 | 0 | 0 | 0 |
| D123 | D125 | 1749 | 6945 | 6933 | 11 | 0 | 1 | 0.16 | 0 | 9.09 |
| D123 | D127 | 1648 | 6363 | 6344 | 19 | 0 | 4 | 0.3 | 0 | 21.05 |
| D123 | D129 | 1675 | 6385 | 6360 | 23 | 2 | 5 | 0.36 | 8.7 | 21.74 |
| D123 | D131 | 1628 | 6379 | 6373 | 4 | 0 | 2 | 0.06 | 0 | 50 |
| D123 | D132 | 1617 | 6421 | 6399 | 18 | 0 | 4 | 0.28 | 0 | 22.22 |
| D123 | D133 | 1745 | 6830 | 6802 | 25 | 0 | 4 | 0.37 | 0 | 16 |
| D123 | D134 | 1729 | 6740 | 6719 | 19 | 0 | 6 | 0.28 | 0 | 31.58 |
| D124 | D018 | 1700 | 6627 | 6597 | 27 | 2 | 7 | 0.41 | 7.41 | 25.93 |
| D124 | D123 | 3057 | 12266 | 12266 | 0 | 0 | 0 | 0 | 0 | 0 |
| D124 | D125 | 1749 | 6942 | 6930 | 11 | 0 | 1 | 0.16 | 0 | 9.09 |
| D124 | D127 | 1648 | 6361 | 6342 | 19 | 0 | 4 | 0.3 | 0 | 21.05 |
| D124 | D129 | 1678 | 6387 | 6362 | 23 | 2 | 5 | 0.36 | 8.7 | 21.74 |
| D124 | D131 | 1631 | 6383 | 6376 | 5 | 0 | 2 | 0.08 | 0 | 40 |
| D124 | D132 | 1617 | 6423 | 6400 | 19 | 0 | 4 | 0.3 | 0 | 21.05 |
| D124 | D133 | 1747 | 6832 | 6805 | 24 | 0 | 4 | 0.35 | 0 | 16.67 |
| D124 | D134 | 1729 | 6739 | 6717 | 20 | 0 | 5 | 0.3 | 0 | 25 |
| D125 | D018 | 1674 | 6578 | 6556 | 19 | 0 | 2 | 0.29 | 0 | 10.53 |
| D125 | D123 | 1749 | 6945 | 6933 | 11 | 0 | 0 | 0.16 | 0 | 0 |
| D125 | D124 | 1749 | 6942 | 6930 | 11 | 0 | 0 | 0.16 | 0 | 0 |
| D125 | D127 | 1671 | 6390 | 6376 | 12 | 0 | 1 | 0.19 | 0 | 8.33 |
| D125 | D129 | 1641 | 6261 | 6240 | 17 | 0 | 2 | 0.27 | 0 | 11.76 |
| D125 | D131 | 1718 | 6707 | 6691 | 12 | 0 | 3 | 0.18 | 0 | 25 |
| D125 | D132 | 1718 | 6640 | 6620 | 15 | 0 | 1 | 0.23 | 0 | 6.67 |
| D125 | D133 | 1700 | 6601 | 6584 | 10 | 0 | 3 | 0.15 | 0 | 30 |
| D125 | D134 | 1677 | 6567 | 6549 | 14 | 0 | 3 | 0.21 | 0 | 21.43 |
| D127 | D018 | 1578 | 6096 | 6067 | 26 | 4 | 3 | 0.43 | 15.38 | 11.54 |
| D127 | D123 | 1648 | 6363 | 6344 | 19 | 0 | 1 | 0.3 | 0 | 5.26 |
| D127 | D124 | 1648 | 6361 | 6342 | 19 | 0 | 2 | 0.3 | 0 | 10.53 |
| D127 | D125 | 1671 | 6390 | 6376 | 12 | 0 | 0 | 0.19 | 0 | 0 |
| D127 | D129 | 1589 | 6000 | 5971 | 26 | 0 | 5 | 0.43 | 0 | 19.23 |
| D127 | D131 | 1652 | 6347 | 6339 | 7 | 0 | 0 | 0.11 | 0 | 0 |
| D127 | D132 | 1670 | 6443 | 6429 | 10 | 0 | 1 | 0.16 | 0 | 10 |
| D127 | D133 | 1585 | 6200 | 6181 | 14 | 0 | 3 | 0.23 | 0 | 21.43 |
| D127 | D134 | 1684 | 6424 | 6404 | 18 | 0 | 4 | 0.28 | 0 | 22.22 |
| D129 | D018 | 1603 | 6164 | 6147 | 15 | 4 | 0 | 0.24 | 26.67 | 0 |
| D129 | D123 | 1675 | 6385 | 6360 | 23 | 2 | 7 | 0.36 | 8.7 | 30.43 |
| D129 | D124 | 1678 | 6387 | 6362 | 23 | 2 | 7 | 0.36 | 8.7 | 30.43 |
| D129 | D125 | 1641 | 6261 | 6240 | 17 | 2 | 5 | 0.27 | 11.76 | 29.41 |
| D129 | D127 | 1589 | 6000 | 5971 | 26 | 2 | 5 | 0.43 | 7.69 | 19.23 |
| D129 | D131 | 1616 | 6066 | 6047 | 18 | 5 | 3 | 0.3 | 27.78 | 16.67 |
| D129 | D132 | 1591 | 6187 | 6160 | 24 | 0 | 2 | 0.39 | 0 | 8.33 |
| D129 | D133 | 1669 | 6717 | 6702 | 15 | 0 | 0 | 0.22 | 0 | 0 |
| D129 | D134 | 1757 | 6647 | 6633 | 12 | 2 | 1 | 0.18 | 16.67 | 8.33 |
| D131 | D018 | 1589 | 6168 | 6149 | 16 | 2 | 3 | 0.26 | 12.5 | 18.75 |
| D131 | D123 | 1628 | 6379 | 6373 | 4 | 0 | 0 | 0.06 | 0 | 0 |
| D131 | D124 | 1631 | 6383 | 6376 | 5 | 0 | 0 | 0.08 | 0 | 0 |

| | | | | | | | | | | |
|---|---|---|---|---|---|---|---|---|---|---|
| D131 | D125 | 1718 | 6707 | 6691 | 12 | 0 | 2 | 0.18 | 0 | 16.67 |
| D131 | D127 | 1652 | 6347 | 6339 | 7 | 0 | 1 | 0.11 | 0 | 14.29 |
| D131 | D129 | 1616 | 6066 | 6047 | 18 | 2 | 3 | 0.3 | 11.11 | 16.67 |
| D131 | D132 | 1618 | 6332 | 6310 | 17 | 2 | 0 | 0.27 | 11.76 | 0 |
| D131 | D133 | 1610 | 6228 | 6212 | 14 | 2 | 1 | 0.22 | 14.29 | 7.14 |
| D131 | D134 | 1689 | 6478 | 6461 | 15 | 0 | 3 | 0.23 | 0 | 20 |
| D132 | D018 | 1611 | 6256 | 6235 | 18 | 4 | 3 | 0.29 | 22.22 | 16.67 |
| D132 | D123 | 1617 | 6421 | 6399 | 18 | 0 | 3 | 0.28 | 0 | 16.67 |
| D132 | D124 | 1617 | 6423 | 6400 | 19 | 0 | 3 | 0.3 | 0 | 15.79 |
| D132 | D125 | 1718 | 6640 | 6620 | 15 | 2 | 3 | 0.23 | 13.33 | 20 |
| D132 | D127 | 1670 | 6443 | 6429 | 10 | 0 | 1 | 0.16 | 0 | 10 |
| D132 | D129 | 1591 | 6187 | 6160 | 24 | 0 | 3 | 0.39 | 0 | 12.5 |
| D132 | D131 | 1618 | 6332 | 6310 | 17 | 5 | 1 | 0.27 | 29.41 | 5.88 |
| D132 | D133 | 1693 | 6837 | 6824 | 9 | 0 | 1 | 0.13 | 0 | 11.11 |
| D132 | D134 | 1644 | 6307 | 6280 | 24 | 4 | 2 | 0.38 | 16.67 | 8.33 |
| D133 | D018 | 1686 | 6582 | 6563 | 16 | 2 | 1 | 0.24 | 12.5 | 6.25 |
| D133 | D123 | 1745 | 6830 | 6802 | 25 | 0 | 3 | 0.37 | 0 | 12 |
| D133 | D124 | 1747 | 6832 | 6805 | 24 | 0 | 3 | 0.35 | 0 | 12.5 |
| D133 | D125 | 1700 | 6601 | 6584 | 10 | 2 | 2 | 0.15 | 20 | 20 |
| D133 | D127 | 1585 | 6200 | 6181 | 14 | 0 | 5 | 0.23 | 0 | 35.71 |
| D133 | D129 | 1669 | 6717 | 6702 | 15 | 0 | 2 | 0.22 | 0 | 13.33 |
| D133 | D131 | 1610 | 6228 | 6212 | 14 | 5 | 0 | 0.22 | 35.71 | 0 |
| D133 | D132 | 1693 | 6837 | 6824 | 9 | 0 | 3 | 0.13 | 0 | 33.33 |
| D133 | D134 | 1736 | 6711 | 6690 | 18 | 4 | 3 | 0.27 | 22.22 | 16.67 |
| D134 | D018 | 1820 | 7040 | 7024 | 14 | 0 | 0 | 0.2 | 0 | 0 |
| D134 | D123 | 1729 | 6740 | 6719 | 19 | 0 | 6 | 0.28 | 0 | 31.58 |
| D134 | D124 | 1729 | 6739 | 6717 | 20 | 0 | 6 | 0.3 | 0 | 30 |
| D134 | D125 | 1677 | 6567 | 6549 | 14 | 0 | 4 | 0.21 | 0 | 28.57 |
| D134 | D127 | 1684 | 6424 | 6404 | 18 | 0 | 6 | 0.28 | 0 | 33.33 |
| D134 | D129 | 1757 | 6647 | 6633 | 12 | 0 | 2 | 0.18 | 0 | 16.67 |
| D134 | D131 | 1689 | 6478 | 6461 | 15 | 0 | 3 | 0.23 | 0 | 20 |
| D134 | D132 | 1644 | 6307 | 6280 | 24 | 2 | 2 | 0.38 | 8.33 | 8.33 |
| D134 | D133 | 1736 | 6711 | 6690 | 18 | 2 | 4 | 0.27 | 11.11 | 22.22 |

**Table S4.** Pairwise comparison of overlapping phased positions in *Aspergillus nidulans* phased isolates, representing a positive control of meiotic crossover identification (OPG = Overlapping Phase Groups, OOP = Overlapping Phased Positions, GC = Gene Conversion).

| Isolate A | Isolate B | OPG | OPP | OPP Same | OPP Cross | OPP Cross (GC) | OPP Cross (Meiosis) | OPP Cross (%) | OPP Cross (GC)% | OPP Cross (Meiosis)% |
|---|---|---|---|---|---|---|---|---|---|---|
| S117 | S118 | 8369 | 39101 | 39101 | 0 | 0 | 0 | 0 | 0 | 0 |
| S117 | S119 | 8374 | 38514 | 38512 | 2 | 0 | 0 | 0.01 | 0 | 0 |
| S117 | S120 | 8367 | 38256 | 38254 | 2 | 0 | 0 | 0.01 | 0 | 0 |
| S117 | S121 | 8403 | 39115 | 39113 | 1 | 0 | 0 | 0 | 0 | 0 |
| S117 | S79 | 7877 | 35767 | 35762 | 5 | 0 | 0 | 0.01 | 0 | 0 |
| S117 | S80 | 7894 | 35616 | 35608 | 7 | 0 | 0 | 0.02 | 0 | 0 |
| S117 | S81 | 7864 | 35929 | 35922 | 5 | 0 | 0 | 0.01 | 0 | 0 |
| S117 | S82 | 7897 | 36032 | 36023 | 6 | 0 | 1 | 0.02 | 0 | 16.67 |
| S117 | S83 | 7829 | 35000 | 34994 | 4 | 0 | 0 | 0.01 | 0 | 0 |
| S118 | S117 | 8369 | 39101 | 39101 | 0 | 0 | 0 | 0 | 0 | 0 |
| S118 | S119 | 8371 | 39024 | 39024 | 0 | 0 | 0 | 0 | 0 | 0 |
| S118 | S120 | 8346 | 38653 | 38651 | 1 | 0 | 0 | 0 | 0 | 0 |
| S118 | S121 | 8580 | 41066 | 41063 | 1 | 0 | 0 | 0 | 0 | 0 |
| S118 | S79 | 7141 | 33025 | 33021 | 3 | 0 | 0 | 0.01 | 0 | 0 |
| S118 | S80 | 7178 | 32944 | 32938 | 5 | 0 | 3 | 0.02 | 0 | 60 |
| S118 | S81 | 7175 | 33295 | 33293 | 1 | 0 | 0 | 0 | 0 | 0 |
| S118 | S82 | 7199 | 33368 | 33360 | 6 | 0 | 3 | 0.02 | 0 | 50 |
| S118 | S83 | 7132 | 32299 | 32293 | 5 | 0 | 2 | 0.02 | 0 | 40 |
| S119 | S117 | 8374 | 38514 | 38512 | 2 | 0 | 0 | 0.01 | 0 | 0 |
| S119 | S118 | 8371 | 39024 | 39024 | 0 | 0 | 0 | 0 | 0 | 0 |
| S119 | S120 | 8377 | 38212 | 38211 | 1 | 0 | 0 | 0 | 0 | 0 |
| S119 | S121 | 8406 | 39060 | 39060 | 0 | 0 | 0 | 0 | 0 | 0 |
| S119 | S79 | 7137 | 32619 | 32616 | 3 | 0 | 0 | 0.01 | 0 | 0 |
| S119 | S80 | 7187 | 32598 | 32592 | 6 | 0 | 3 | 0.02 | 0 | 50 |
| S119 | S81 | 7174 | 32902 | 32898 | 3 | 0 | 0 | 0.01 | 0 | 0 |
| S119 | S82 | 7198 | 32949 | 32943 | 5 | 0 | 2 | 0.02 | 0 | 40 |
| S119 | S83 | 7169 | 32087 | 32085 | 1 | 0 | 0 | 0 | 0 | 0 |
| S120 | S117 | 8367 | 38256 | 38254 | 2 | 0 | 0 | 0.01 | 0 | 0 |
| S120 | S118 | 8346 | 38653 | 38651 | 1 | 0 | 0 | 0 | 0 | 0 |
| S120 | S119 | 8377 | 38212 | 38211 | 1 | 0 | 0 | 0 | 0 | 0 |
| S120 | S121 | 8376 | 38678 | 38676 | 1 | 0 | 0 | 0 | 0 | 0 |
| S120 | S79 | 7162 | 32527 | 32522 | 4 | 0 | 1 | 0.01 | 0 | 25 |
| S120 | S80 | 7202 | 32483 | 32476 | 6 | 0 | 2 | 0.02 | 0 | 33.33 |
| S120 | S81 | 7173 | 32732 | 32729 | 2 | 0 | 1 | 0.01 | 0 | 50 |
| S120 | S82 | 7158 | 32723 | 32717 | 5 | 0 | 2 | 0.02 | 0 | 40 |
| S120 | S83 | 7123 | 31816 | 31813 | 2 | 0 | 0 | 0.01 | 0 | 0 |
| S121 | S117 | 8403 | 39115 | 39113 | 1 | 0 | 0 | 0 | 0 | 0 |
| S121 | S118 | 8580 | 41066 | 41063 | 1 | 0 | 0 | 0 | 0 | 0 |
| S121 | S119 | 8406 | 39060 | 39060 | 0 | 0 | 0 | 0 | 0 | 0 |
| S121 | S120 | 8376 | 38678 | 38676 | 1 | 0 | 0 | 0 | 0 | 0 |
| S121 | S79 | 7142 | 33028 | 33024 | 4 | 0 | 0 | 0.01 | 0 | 0 |
| S121 | S80 | 7194 | 32885 | 32879 | 6 | 0 | 3 | 0.02 | 0 | 50 |
| S121 | S81 | 7193 | 33291 | 33287 | 2 | 0 | 0 | 0.01 | 0 | 0 |
| S121 | S82 | 7224 | 33372 | 33361 | 8 | 0 | 2 | 0.02 | 0 | 25 |
| S121 | S83 | 7141 | 32307 | 32302 | 4 | 0 | 2 | 0.01 | 0 | 50 |
| S79 | S117 | 7877 | 35767 | 35762 | 5 | 0 | 0 | 0.01 | 0 | 0 |
| S79 | S118 | 7141 | 33025 | 33021 | 3 | 0 | 0 | 0.01 | 0 | 0 |
| S79 | S119 | 7137 | 32619 | 32616 | 3 | 0 | 0 | 0.01 | 0 | 0 |
| S79 | S120 | 7162 | 32527 | 32522 | 4 | 0 | 0 | 0.01 | 0 | 0 |
| S79 | S121 | 7142 | 33028 | 33024 | 4 | 0 | 1 | 0.01 | 0 | 25 |
| S79 | S80 | 9715 | 43241 | 43239 | 2 | 0 | 0 | 0 | 0 | 0 |
| S79 | S81 | 9693 | 43662 | 43661 | 1 | 0 | 0 | 0 | 0 | 0 |
| S79 | S82 | 9724 | 43666 | 43665 | 1 | 0 | 0 | 0 | 0 | 0 |
| S79 | S83 | 9623 | 42473 | 42471 | 2 | 0 | 1 | 0 | 0 | 50 |
| S80 | S117 | 7894 | 35616 | 35608 | 7 | 0 | 0 | 0.02 | 0 | 0 |
| S80 | S118 | 7178 | 32944 | 32938 | 5 | 0 | 1 | 0.02 | 0 | 20 |
| S80 | S119 | 7187 | 32598 | 32592 | 6 | 0 | 0 | 0.02 | 0 | 0 |

| | | | | | | | | | | |
|---|---|---|---|---|---|---|---|---|---|---|
| S80 | S120 | 7202 | 32483 | 32476 | 6 | 0 | 1 | 0.02 | 0 | 16.67 |
| S80 | S121 | 7194 | 32885 | 32879 | 6 | 0 | 1 | 0.02 | 0 | 16.67 |
| S80 | S79 | 9715 | 43241 | 43239 | 2 | 0 | 1 | 0 | 0 | 50 |
| S80 | S81 | 9722 | 43496 | 43494 | 1 | 0 | 0 | 0 | 0 | 0 |
| S80 | S82 | 9740 | 43523 | 43518 | 4 | 0 | 1 | 0.01 | 0 | 25 |
| S80 | S83 | 9660 | 42371 | 42370 | 1 | 0 | 0 | 0 | 0 | 0 |
| S81 | S117 | 7864 | 35929 | 35922 | 5 | 0 | 0 | 0.01 | 0 | 0 |
| S81 | S118 | 7175 | 33295 | 33293 | 1 | 0 | 0 | 0 | 0 | 0 |
| S81 | S119 | 7174 | 32902 | 32898 | 3 | 0 | 0 | 0.01 | 0 | 0 |
| S81 | S120 | 7173 | 32732 | 32729 | 2 | 0 | 0 | 0.01 | 0 | 0 |
| S81 | S121 | 7193 | 33291 | 33287 | 2 | 0 | 0 | 0.01 | 0 | 0 |
| S81 | S79 | 9693 | 43662 | 43661 | 1 | 0 | 1 | 0 | 0 | 100 |
| S81 | S80 | 9722 | 43496 | 43494 | 1 | 0 | 0 | 0 | 0 | 0 |
| S81 | S82 | 9735 | 43976 | 43974 | 2 | 0 | 0 | 0 | 0 | 0 |
| S81 | S83 | 9668 | 42776 | 42775 | 1 | 0 | 0 | 0 | 0 | 0 |
| S82 | S117 | 7897 | 36032 | 36023 | 6 | 0 | 0 | 0.02 | 0 | 0 |
| S82 | S118 | 7199 | 33368 | 33360 | 6 | 0 | 2 | 0.02 | 0 | 33.33 |
| S82 | S119 | 7198 | 32949 | 32943 | 5 | 0 | 2 | 0.02 | 0 | 40 |
| S82 | S120 | 7158 | 32723 | 32717 | 5 | 0 | 0 | 0.02 | 0 | 0 |
| S82 | S121 | 7224 | 33372 | 33361 | 8 | 0 | 1 | 0.02 | 0 | 12.5 |
| S82 | S79 | 9724 | 43666 | 43665 | 1 | 0 | 0 | 0 | 0 | 0 |
| S82 | S80 | 9740 | 43523 | 43518 | 4 | 0 | 0 | 0.01 | 0 | 0 |
| S82 | S81 | 9735 | 43976 | 43974 | 2 | 0 | 0 | 0 | 0 | 0 |
| S82 | S83 | 9682 | 42808 | 42807 | 0 | 0 | 0 | 0 | 0 | 0 |
| S83 | S117 | 7829 | 35000 | 34994 | 4 | 0 | 0 | 0.01 | 0 | 0 |
| S83 | S118 | 7132 | 32299 | 32293 | 5 | 0 | 2 | 0.02 | 0 | 40 |
| S83 | S119 | 7169 | 32087 | 32085 | 1 | 0 | 0 | 0 | 0 | 0 |
| S83 | S120 | 7123 | 31816 | 31813 | 2 | 0 | 1 | 0.01 | 0 | 50 |
| S83 | S121 | 7141 | 32307 | 32302 | 4 | 0 | 1 | 0.01 | 0 | 25 |
| S83 | S79 | 9623 | 42473 | 42471 | 2 | 0 | 1 | 0 | 0 | 50 |
| S83 | S80 | 9660 | 42371 | 42370 | 1 | 0 | 0 | 0 | 0 | 0 |
| S83 | S81 | 9668 | 42776 | 42775 | 1 | 0 | 0 | 0 | 0 | 0 |
| S83 | S82 | 9682 | 42808 | 42807 | 0 | 0 | 0 | 0 | 0 | 0 |

**Table S5.** Pairwise comparison of overlapping phased positions in *C. albicans* phased isolates, representing a negative control of meiotic crossover identification (OPG = Overlapping Phase Groups, OOP = Overlapping Phased Positions, GC = Gene Conversion).

| Isolate A | Isolate B | OPG | OPP | OPP Same | OPP Cross | OPP Cross (GC) | OPP Cross (Meiosis) | OPP Cross (%) | OPP Cross (GC)% | OPP Cross (Meiosis)% |
|---|---|---|---|---|---|---|---|---|---|---|
| 881205 | ATCC48184 | 29344 | 395843 | 395827 | 11 | 0 | 3 | 0.00 | 0.00 | 27.27 |
| 881205 | CBS132 | 21539 | 302628 | 299139 | 221 | 173 | 14 | 0.07 | 78.28 | 6.33 |
| 881205 | CBS7821 | 38324 | 517748 | 511577 | 393 | 322 | 21 | 0.08 | 81.93 | 5.34 |
| ATCC48184 | 881205 | 29344 | 395843 | 395827 | 11 | 0 | 3 | 0.00 | 0.00 | 27.27 |
| ATCC48184 | CBS132 | 19128 | 241575 | 238828 | 165 | 124 | 14 | 0.07 | 75.15 | 8.48 |
| ATCC48184 | CBS7821 | 30897 | 376272 | 371777 | 282 | 246 | 8 | 0.08 | 87.23 | 2.84 |
| CBS132 | 881205 | 21539 | 302628 | 299139 | 221 | 173 | 16 | 0.07 | 78.28 | 7.24 |
| CBS132 | ATCC48184 | 19128 | 241575 | 238828 | 165 | 124 | 11 | 0.07 | 75.15 | 6.67 |
| CBS132 | CBS7821 | 23442 | 329310 | 328006 | 47 | 28 | 6 | 0.01 | 59.57 | 12.77 |
| CBS7821 | 881205 | 38324 | 517748 | 511577 | 393 | 322 | 17 | 0.08 | 81.93 | 4.33 |
| CBS7821 | ATCC48184 | 30897 | 376272 | 371777 | 282 | 246 | 8 | 0.08 | 87.23 | 2.84 |
| CBS7821 | CBS132 | 23442 | 329310 | 328006 | 47 | 28 | 5 | 0.01 | 59.57 | 10.64 |

**Table S6.** Pairwise comparison of overlapping phased positions in *C. neoformans* hybrid isolates showing crossover counts within the genome

#### Supplementary Methods

Ortholog prediction and synteny analysis were performed using Synima v2.0.0 (2). Orthologs were inferred using orthomcl v1.4 (3) based on an all-vs-all comparison of peptide sequences computed with diamond v2.1.6.160 (4) using the parameters max_target_seqs=250, evalue=1e-10, diamond_sensitivity=fast. Orthogroups assigned by Synima were classified into core, accessory, and unique categories. 2060 single-copy core orthologs were identified and used to construct a phylogenetic tree. Each orthogroup of single-copy orthologs was aligned separately using MUSCLE v. 5.3.osx64 (5) with default settings. All alignments were concatenated into a single FASTA, and an 'approximately maximum-likelihood' tree was inferred using FastTree v2.1.11 SSE3 (6). Synteny blocks were identified as chains of ≥ 4 orthologous genes using DAGChainer (7), and visualised using Synima.
